# The Cost to Scroll: Short-form Video Use and Episodic Memory

**DOI:** 10.64898/2026.10.07.757368

**Authors:** Kevin K. Chen, Douglas S. Miller, Khanh K. Nguyen, Anthony D. Wagner, Tammy T. Tran

## Abstract

Short-form video platforms expose users to rapid streams of novel content; these features are posited to place heavy demands on cognitive processes. In a preregistered online study, we used device-reported screen time (TikTok and total Instagram app use) to examine whether short-form video use is associated with sustained attention and episodic memory in participants aged 18–24 yrs (*N* = 497). During a sustained attention task, participants made indoor/outdoor decisions on scene stimuli, with 90% frequent trials (indoor images) and 10% infrequent trials (outdoor images); the subsequent surprise recognition test assessed memory for these items. Among participants with device-reported TikTok or Instagram use (*N* = 395), short-form video use was not significantly associated with attention performance. By contrast, greater short-form video use was associated with poorer episodic memory (*β* = *−*0.15, *p* =.004); this relationship was strongest for infrequent-category items. These observational findings connect heavier short-form video consumption with poorer episodic memory in young adults, showing how daily digital habits co-vary with memory function.

## 1 Introduction

Short-form video has emerged as a dominant mode of daily digital consumption, particularly among young adults, by delivering highly personalized, algorithmically curated streams of content^1,2^. Short-form video formats are now widespread across major digital platforms, including TikTok, Instagram Reels and YouTube Shorts^1^. Recent demographic data indicate that social media platform use is widespread among young adults, with adults under 30 yrs especially likely to use platforms such as Instagram, Snapchat, TikTok and Reddit^3^. Yet, despite widespread public and academic concern regarding the potential adverse effects of intensive social media consumption on human cognition, empirical research directly linking short-form video use behavior to cognitive differences remains mixed.

Within the attention domain, a recent meta-analysis found that greater short-form video use was associated with poorer attention ^1^. For instance, greater self-reported short-form video use was associated with lower attentional control among adolescents ^4^, and individuals classified as addicted short-form video users exhibited poorer attentional performance during a video-watching task and on a subsequent Stroop task ^5^. In a separate study, heavy short-form social media users who reported more than 4 hrs of daily engagement showed an attenuated P300 response during a visual oddball paradigm relative to lower-frequency users who reported less than 3 hrs of daily engagement^6^. In the survey phase of Lin et al., ^7^ greater self-reported short-form video use was associated with poorer sustained-attention performance, including increased commission errors and reaction-time variability on a sustained attention to response task.

On the other hand, negative relationships between social media use and attention are not consistently observed. Findings from studies using objective behavioral measures of attention are also mixed. An attentional-bias study found that attentional bias toward social media stimuli was not associated with social media engagement or addiction-severity measures and found no evidence of attentional bias toward those stimuli^8^. Work with objective device-logged measures found only small effects, with greater social media and TikTok use associated with slightly better, rather than worse, attentional control ^9^.

Given that the core architectural model of short-form video relies on rapid, highly engaging events composed of high-frequency content shifts, episodic memory function may also vary with short-form video use. While little is known about this possible relationship, prior studies have found poorer prospective memory and visual short-term memory following short-form video exposure ^10,11^, raising the possibility that short-form video exposure interferes with both the maintenance of mnemonic goals and the brief retention of visual information. Yet, a fundamental question remains: are mnemonic impairments transient, acute effects that immediately follow social media usage, or are there more enduring differences in mnemonic processing that relate to short-form video use patterns and do they extend to episodic memory?

Beyond uncertainty about whether observed relationships between short-form video use and cognition are acute or enduring, extant studies are often limited by small samples and reliance on retrospective self-report measures of short-form video use, which may diverge from app-or device-based media-use reports ^9,12^. Notably, Pablo et al. ^12^ used verified media usage data from participants’ devices and found no relationship between media usage and working-memory performance. These findings underscore the importance of distinguishing retrospective self-report estimates from objective app-and device-based estimates when characterizing relationships between short-form video use and cognitive performance.

To address these open questions, we conducted a preregistered study (https://osf.io/3vp7f/ overview?view_only=6608748ac569427d8fa16df974696c3e) examining relationships among short-form video use, sustained attention and episodic memory in young adults. Participants (*N* = 497) were recruited via Prolific, a crowdsourced participant platform, and completed an adapted variant of the Sustained Attention to Response Task (SART) that included a subsequent surprise episodic memory test (SART-M) examining recognition memory for the stimuli encountered during the SART phase^13^. This paradigm allowed us to quantify sustained attention and episodic memory for incidentally encoded stimuli within the same experimental session. Alongside self-reported platform use, we quantified TikTok and Instagram use from smartphone screen-time screenshots. Our primary analyses tested whether greater device-reported aggregate short-form video use was associated with poorer sustained attention and episodic memory. We also examined addiction-like tendencies via the TikTok Addiction Scale (TAS)^14^, media multitasking via the Media Multitasking Index (MMI) ^15^, and impulsivity via the Barratt Impulsiveness Scale (BIS-11)^16^. The present design was motivated by an earlier local-sample pilot version of the task (*N* = 338), composed primarily of Stanford University students and Bay Area young adults. The pilot study used retrospective self-report measures of short-form video use; results from this earlier study were broadly similar to those reported in the present study (see Supplementary Tables S1 and S2 and Supplementary Fig. S1 for participant characteristics and primary continuous associations from the pilot study).

## 2 Methods

### 2.1 Participants

Participants were recruited through Prolific ^17^ between July 2025 and March 2026. Eligibility criteria required participants to be 18–24 yrs old, US-based, fluent in English, have normal or corrected-to-normal vision, have a Prolific approval rate of at least 95%, and complete the study on a desktop or laptop computer using a physical keyboard. All participants provided informed consent before beginning the study. Sample size was determined a priori based on variability observed in the pilot data, in which approximately one-third of participants were non-TikTok users, as well as anticipated participant dropout and data-quality loss on Prolific. We preregistered a plan to recruit 550 participants with the goal of retaining approximately 500 participants, including approximately 200 non-TikTok users and 300 TikTok users. Recruitment was extended to obtain sufficient usable data. Usage–cognition associations were not examined until data collection was complete. A retrospective sensitivity analysis indicated that the primary attention *d^′^*model among device-reported aggregate-use users (*N* = 395) had 80% power to detect an absolute partial correlation of *r* = 0.14, controlling for sex, at a two-sided *α* =.05.

### 2.2 Exclusion criteria and participant flow

Exclusion criteria were applied sequentially. Two preliminary test sessions were excluded from all analyses (*N* = 2). Participants were removed if they did not complete the media usage survey (*N* = 3), had no valid SART file (*N* = 17), or had no scorable SART-M file (*N* = 19). For repeat attempts, we selected the first main-task attempt with a recorded image presentation; practice-only restarts were disregarded. Participants with more than one main encoding attempt before their first memory test were excluded, including partial encoding restarts (*N* = 14). The first memory test was required to contain 200 recorded responses; participants with an incomplete first memory test were excluded (*N* = 3), and later retests were not substituted. Later memory repeats did not invalidate an otherwise eligible first test. Data from separate attempts were never combined. For participant-level attention summaries and the low-valid-trial screen, SART trials with a reaction time faster than 200 ms were treated as invalid. Next, we excluded data from participants with a low SART trial count (*<* 80% valid trials; *N* = 9), low attention accuracy (mean frequent-and infrequent-trial accuracy below *M −* 1.5 SD, i.e., 0.731; *N* = 40), high omission rate (greater than Q3 + 3 times IQR, i.e., 0.076; *N* = 17), or negative overall memory *d^′^* (*N* = 16). Due to concerns about outliers in short-form video usage, 13 additional participants with aggregate use more than 1.5*×*IQR above Q3 were removed. An overview of data exclusion criteria is shown in Fig. 1. In total, 153 participants were excluded from the 650 approved Prolific participants, leaving a final analytic sample of 497 participants (Fig. 1). We also evaluated the robustness of the primary findings to alternative cutoffs of 2.0, 2.5 and 3.0 SDs below the mean attention accuracy; these sensitivity analyses produced the same qualitative pattern of results (Supplementary Table S3). The low-valid-trial criterion excluded participants for whom more than 20% of the 500 SART trials were invalid (no response, missing reaction time, or reaction time *<* 200 ms). This broad data-quality screen is distinct from the high-omission-rate criterion, which identified a small number of remaining participants with unusually high non-response among their eligible trials. Different outlier rules were applied to different variables because their distributions and data-quality implications differed: 1.5*×*IQR for aggregate usage (to reduce leverage from extreme values), Q3 + 3*×*IQR for omission rate (because omission distributions are right-skewed), and a mean-based SD threshold for attention accuracy (to flag clear low performers).

**Figure 1:**
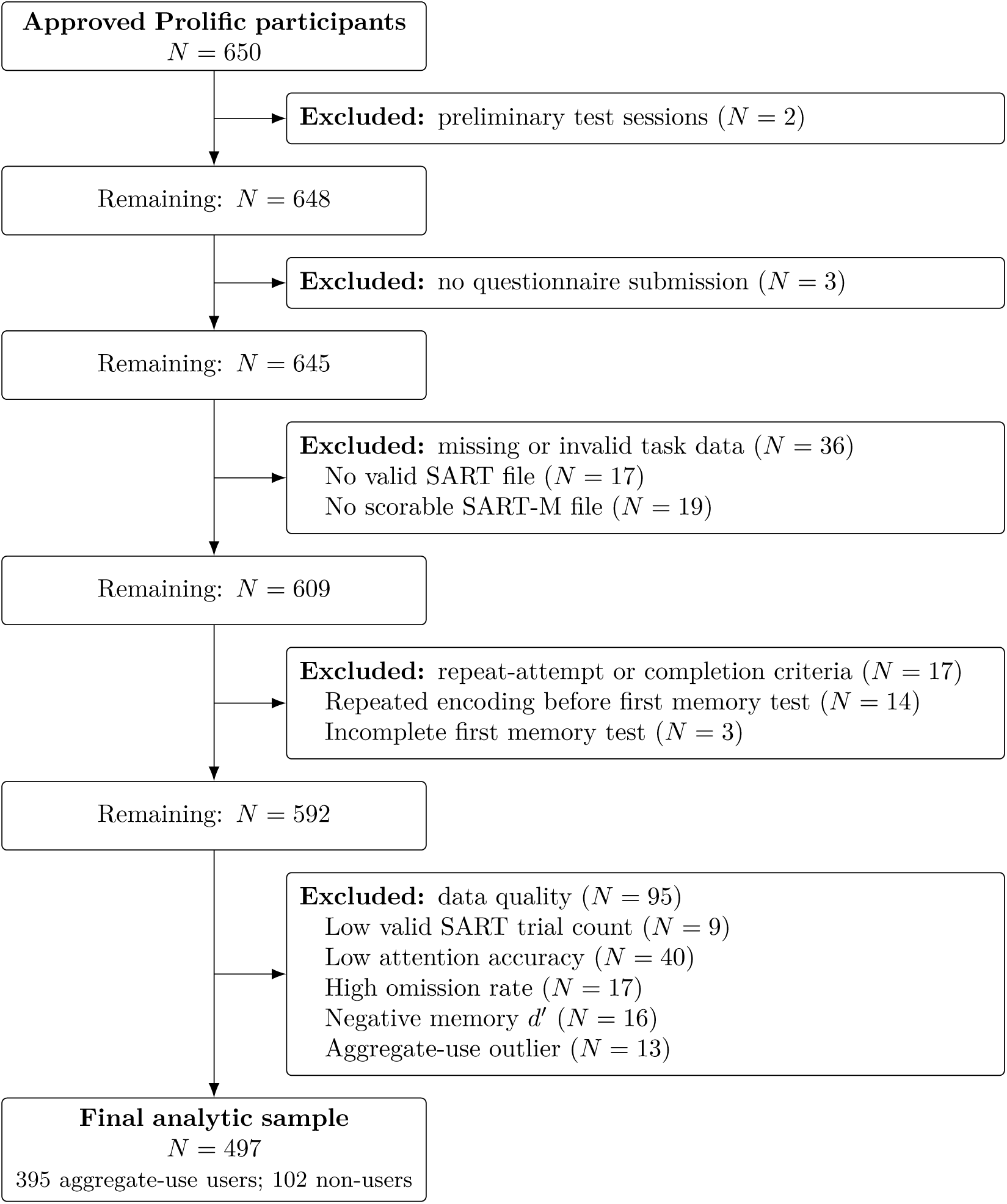
Participant flow diagram. Exclusions were applied sequentially. Boxes on the left show participants remaining after each step; boxes on the right show participants excluded at that step. Exclusion definitions and thresholds are given in Section 2.2. SART, Sustained Attention to Response Task; SART-M, Sustained Attention to Response Task with Memory.

### 2.3 Preregistration

The primary hypotheses and analysis plan were preregistered on the Open Science Framework on July 11, 2025, before collection of the analyzed sample (https://osf.io/3vp7f/overview?view_ only=6608748ac569427d8fa16df974696c3e). Two preliminary test sessions completed on July 10, 2025 were excluded from all analyses. The registration focused on TikTok; the aggregate TikTok and Instagram predictor was added post hoc. Other post hoc additions included condition *×* usage and trial-level models, effect-size reporting, Bayesian and preceding-RT/subsequent-memory analyses, sensitivity analyses and retrospective power calculations. Data-quality thresholds were not preregistered, but followed our local-sample approach and were applied before testing the focal associations. Repeat-attempt handling and the complete-first-memory requirement were finalized after the initial analyses. In the category-specific memory hypothesis, “frequent” should read “infrequent”, consistent with the parallel registered attention and reaction-time hypotheses and an earlier thesis completed before preregistration ^18^ pp. 36, 45. The present report focuses on weekly use, sustained attention and recognition memory; registered past-24-hour, median-split, task-time-course, confidence/ROC, image-memorability and additional questionnaire/composite analyses are not reported here.

### 2.4 Stimuli

Stimuli consisted of high-resolution naturalistic indoor and outdoor scene images drawn from the SUN database^19^. Images were used in both the sustained attention task and the surprise recognition memory test.

During the sustained-attention phase, participants viewed 500 scene images. The task used an imbalanced category structure, with 450 images from the frequent category and 50 images from the infrequent category. In the present implementation, indoor scenes served as the frequent category and outdoor scenes served as the infrequent category. This 9:1 ratio was used to encourage habitual responding and increase sensitivity to attentional lapses, following prior SART-M work ^13^.

The surprise recognition memory test contained 200 trials: 100 old images from the sustained-attention phase and 100 novel foil images. The old images included 50 frequent-category images sampled from the 450 frequent trials and all 50 infrequent-category images. The foil set comprised 50 indoor and 50 outdoor images randomly selected from the SUN database that had not appeared during the sustained-attention phase. Candidate foils judged to be highly visually similar to a studied image were replaced with another randomly selected image based on experimenter perception (K.K.C).

### 2.5 Procedure

The study was conducted online. After informed consent, participants completed the SART-M task and self-report measures assessing short-form video use and individual differences. The full session lasted approximately 30 minutes. For clarity, we refer to the sustained-attention phase as the SART and to the combined sustained-attention and surprise-memory paradigm as the SART-M.

The SART-M task consisted of two phases. First, participants completed a sustained-attention task adapted from the SART-M paradigm^13^. On each trial, participants viewed a scene image for 1000 ms (Fig. 2**A**, upper) and made an indoor/outdoor category judgment using two keyboard keys, “h” or “j.” Only the mapping of response keys to the indoor and outdoor categories was counterbalanced across participants. A central fixation dot was displayed with each image and changed from white to black after a response.

**Figure 2:**
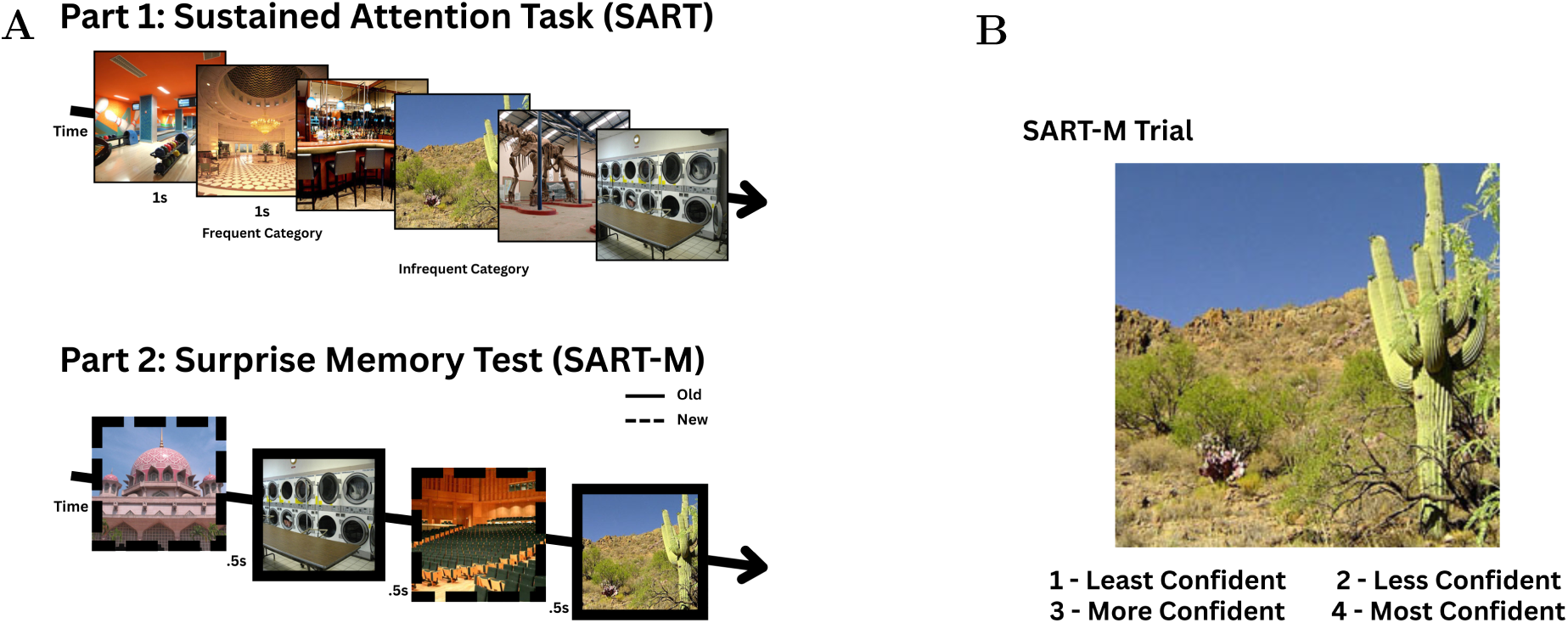
Task procedure and recognition-memory interface. **A**, SART and SART-M task structure, including the sustained-attention phase and subsequent surprise recognition memory test. **B**, Example recognition-memory trial. Participants rated their confidence that the image was old using a 4-point scale.

Immediately after the sustained-attention task, participants completed a surprise recognition memory test for the scenes (Fig. 2**A**, lower). Participants were not told in advance that their memory for the scenes would be tested, ensuring that encoding during the SART phase was incidental. On each memory trial, participants viewed either a studied or novel image and rated their confidence that it had appeared in the earlier task using a 4-point scale, ranging from 1 (lowest confidence that the image was old) to 4 (highest confidence that the image was old). The image and response mappings remained on screen until response (Fig. 2**B**). After a response, the selected response was shown for 500 ms, followed by a 500-ms blank interstimulus interval. The memory task was self-paced.

### 2.6 Sustained-attention outcomes

Sustained-attention performance was quantified using trial-level accuracy and participant-level summary metrics. The primary attention outcomes were attention *d^′^*and condition-specific accuracy for frequent and infrequent trials (incorrect-response rate and omission rate are reported as supplementary descriptive metrics only; Supplementary Fig. S4). Following the preregistered scoring scheme, attention *d^′^*indexed participants’ ability to discriminate frequent and infrequent trial types, calculated as *z*(hit rate) *− z*(false alarm rate), where hit rate was the proportion of responded frequent trials classified correctly and false alarm rate was the proportion of responded infrequent trials classified incorrectly. For all *d^′^* calculations, hit and false-alarm rates of 0 or 1 were adjusted using a 0.5/*N* boundary correction before the inverse-normal transformation, where *N* was the number of trials contributing to the relevant rate. Because attention *d^′^* is defined jointly from the frequent-trial hit rate and infrequent-trial false-alarm rate, separate condition-specific *d^′^* values are not defined for the frequent and infrequent trial categories; condition-specific performance was therefore summarized using accuracy. Incorrect-response rate was calculated as the proportion of responded frequent trials with an incorrect response. Omission rate was calculated separately as the proportion of eligible trials with no recorded response (i.e., a missing or None response key). For attention *d^′^*, condition-specific accuracy and reaction-time summary metrics, trials with no recorded response, a missing reaction time, or a reaction time faster than 200 ms were excluded.

### 2.7 Memory outcomes

Memory performance was assessed using recognition accuracy and signal-detection sensitivity. Recognition responses were scored from the 4-point confidence scale. For images that had appeared during the sustained-attention task, responses of 3 or 4 were scored as correct “old” responses (i.e., hits) and responses of 1 or 2 were scored as incorrect “new” responses (i.e., misses). For foil images that had not appeared during the sustained-attention task, responses of 1 or 2 were scored as correct “new” responses (i.e., correct rejections) and responses of 3 or 4 were scored as false alarms. Overall memory *d^′^* indexed recognition discrimination between old and new items, calculated as *z*(hit rate) *− z*(false alarm rate), where hit rate was the proportion of old images correctly classified as “old” and false alarm rate was the proportion of new foil images incorrectly classified as “old.” Condition-specific memory *d^′^* was computed separately for frequent-category and infrequent-category items, using only the corresponding subset of old images and their matched foils. The same 0.5/*N* boundary correction was applied to extreme hit and false-alarm rates for all memory *d^′^* values. Participant-level memory analyses focused on overall and condition-specific memory *d^′^*. Trial-level models used recognition accuracy, coded according to whether each old or foil image was correctly classified. For trial-level memory models, both studied and foil images were assigned to the frequent or infrequent category according to scene category (indoor vs. outdoor), corresponding to the categories that were frequent or infrequent during encoding.

### 2.8 Short-form video use measures

Short-form video use was assessed for TikTok, Instagram Reels, YouTube Shorts and Facebook Reels, with an additional option for participants to report other short-form video platforms. For each platform, participants provided a retrospective self-report estimate of typical daily use, which was converted to weekly minutes for analysis; free-text entries with explicit hour or minute units were normalized to minutes and ambiguous range entries were treated as missing. For TikTok and Instagram, participants also provided an app-or device-based screenshot report of use over the previous week. Participants entered their usage directly from the relevant app-or device-based screen-time report and uploaded a screenshot as supporting documentation, following prior work using objective mobile screen-time reports to measure social media use^9^. All reported TikTok and Instagram usage values were checked against the uploaded screenshots; when a screenshot was missing or did not match the entered values, participants resubmitted corrected values through a follow-up form, and the corrected values were used. Additional documented corrections were made from the uploaded screenshots: verified weekly totals were used where available, and where only a valid daily-average report was available, weekly use was estimated as daily minutes *×* 7; these source reports did not uniformly cover seven complete days (sensitivity to the daily-average estimates is reported in the Supplementary Information). As these app/device-based reports are less reliant on retrospective estimation than self-reported daily-use estimates, they were used as the primary usage measures for the primary inferential analyses. As such, the primary predictor was device-reported aggregate short-form video use (aggregate use), defined as TikTok minutes plus Instagram minutes over the previous week. The primary predictor was restricted to TikTok and Instagram because device-reported measures were collected only for these platforms; YouTube Shorts and Facebook Reels were assessed by self-report only. As the Instagram screen-time report reflects total app use, this measure includes non-Reels activity (e.g., Stories, feed browsing). Aggregate use therefore serves as a proxy for short-form video engagement. Participants reporting app-or device-based measures of 0 minutes on both TikTok and Instagram were classified as “aggregate-use non-users” for the primary analyses. Participants with non-zero device-reported aggregate use were classified as “aggregate-use users” and were divided into low, medium and high usage tertiles.

### 2.9 Questionnaires

TikTok addiction-like engagement was measured using the TikTok Addiction Scale (TAS) ^14^, an 18-item scale in which responses are summed to produce a total score (TikTok users included in the TAS analysis: *N* = 289, *M* = 40.6, SD = 14.6). Media multitasking was measured using an adapted, truncated version of the Media Multitasking Index (MMI), following the weighted formula introduced by Ophir et al. and used in subsequent work on media multitasking and memory ^15,20^. Participants reported weekly hrs across nine media categories and indicated how often they engaged in each possible pair of media activities concurrently. Responses to the latter were coded as Most of the time = 1, Some of the time = 0.67, A little of the time = 0.33 and Never = 0. As the survey presented each unordered media pair once, every pairwise response was entered symmetrically so that it contributed to the multitasking scores of both constituent media. For each participant, the adapted MMI was calculated as the sum of each medium’s weekly hrs multiplied by its reconstructed multitasking score, divided by total weekly media hrs. One participant had an incomplete pairwise response and was assigned a missing MMI value; among participants with complete MMI data (*N* = 496), *M* = 3.00 (*SD* = 1.29). As this version used fewer media categories than the original Ophir MMI, we refer to it as an adapted/truncated MMI. Impulsivity was measured using the BIS-11^16^, a 30-item scale with a 4-point response format ranging from Rarely/Never to Almost Always/Always and 11 reverse-scored items (full sample: *N* = 497, *M* = 63.9, SD = 11.3). Additional collected demographics included age, sex, race and yrs of education completed.

### 2.10 Statistical analysis

Primary participant-level analyses used multiple linear regression analysis to predict sustained-attention outcomes and memory *d^′^* from aggregate use, with sex included as a covariate. At the trial level, linear mixed-effects models treated attention and recognition-memory accuracy as binary outcomes (correct = 1, incorrect = 0) and estimated the effects of condition, aggregate use, sex, and the condition *×* usage and condition *×* sex interactions, with a random intercept for participant to account for non-independence of repeated trials. Because participant-level covariates otherwise cancel from within-participant condition differences, condition *×* covariate terms were included so that condition *×* usage interactions were adjusted for sex. Each trial-level mixed model was fit with two optimizers, and convergence, a nonzero random-intercept variance and agreement of the estimates across optimizers were verified. The trial-level attention model retained all 500 trials per participant, with no-response trials coded as incorrect, and did not exclude trials on the basis of reaction time; this differs from the participant-level attention summaries, which excluded no-response, missing-reaction-time and *<* 200-ms trials before computing accuracy and *d^′^*.

An exploratory Bayesian multiple linear regression additionally estimated the association between aggregate use and attention *d^′^* among aggregate-use users, controlling for sex. Continuous variables were standardized before model fitting in brms^21^; weakly informative Normal(0,1) priors were used for regression coefficients and Student-t(3,0,2.5) priors for the intercept and residual standard deviation.

To formally test whether the association between aggregate use and memory differed between frequent-and infrequent-category items, we additionally fit participant-level repeated-measures linear mixed-effects models among aggregate-use users. Condition-specific memory *d^′^* values were entered in long format, with memory condition (frequent vs. infrequent), standardized aggregate use, sex, and the condition *×* use and condition *×* sex interactions as fixed effects and participant as a random intercept. The focal effect was the condition *×* usage interaction. Aggregate use was standardized across aggregate-use users, whereas memory *d^′^* remained on its original scale; interaction coefficients (*b*) therefore reflect the change in the difference between infrequent-and frequent-category memory *d^′^*associated with a one-SD increase in aggregate use.

To test whether the association between aggregate use and memory remained after accounting for sustained-attention performance, we fit adjusted participant-level regression models among aggregate-use users. The model for overall memory *d^′^* included overall attention *d^′^*, aggregate use and sex. These models follow the registration’s narrative description of attention adjustment; the registered formula instead listed RT variability. Because condition-specific attention *d^′^* is undefined (see Sustained-attention outcomes), models for frequent-and infrequent-category memory *d^′^* instead controlled for frequent-trial and infrequent-trial attention accuracy, respectively, together with aggregate use and sex. We also fit a formal attention-adjusted participant-level repeated-measures model that extended the model described above by including overall attention *d^′^* and the condition *×* attention *d^′^* interaction.

For the exploratory deBettencourt-style analysis^13^, we estimated an unpenalized binomial logistic model within each participant and then treated participants as the independent sampling units. Group inference used 100,000 bootstrap resamples of the 496 estimable participant slopes, sampled with replacement; each iteration’s statistic was the mean slope. The 95% CI was the 2.5th–97.5th percentile interval of these bootstrap means. The original-style directional one-sided *p* value was the proportion of bootstrap means less than or equal to zero; the two-sided *p* value was twice the smaller of the proportions at or below zero and at or above zero, capped at 1. These procedures were specified after data collection and were not preregistered.

Secondary participant-level models compared aggregate-use users and non-users on attention and memory outcomes, controlling for sex. Usage-tertile models compared Low, Medium and High aggregate-use groups, with sex included as a covariate. Omnibus tertile effects were evaluated with *F* tests, and pairwise comparisons were corrected using the Holm procedure. Except where explicitly corrected, exploratory analyses report unadjusted *p* values. Tertile figures show sex-adjusted estimated marginal means with model-based 95% confidence intervals, which are appropriate for between-group comparisons; differences between tertiles were evaluated with the Holm-corrected tests rather than by interval overlap.

For analyses of the preregistered Prolific sample, participant-level OLS models with continuous predictors and outcomes are reported as standardized regression coefficients (*β*), with continuous variables *z*-scored before model fitting. For both the attention and memory trials, trial-level linear mixed-effects models used a binary score on each trial (1 = correct, 0 = incorrect). As accuracy outcomes and categorical predictors were not standardized, trial-level effects are summarized using unstandardized accuracy-scale coefficients (*b*), standard errors, 95% confidence intervals and two-sided *p* values. Participant-level repeated-measures memory models used condition-specific memory *d^′^* on its original scale with standardized aggregate use; these interaction effects are therefore reported as unstandardized coefficients (*b*) in memory *d^′^*units per one-SD increase in aggregate use. For significant participant-level continuous associations highlighted in the primary and follow-up analyses, 95% confidence intervals and Cohen’s *d* are also reported. Adapting the continuous-exposure framework of Mathur and VanderWeele, ^22^ Cohen’s *d* was expressed as the standardized outcome difference corresponding to a two-standard-deviation contrast in the predictor (i.e., one standard deviation below to one standard deviation above the mean); for covariate-adjusted models the residual standard deviation of the fitted model was used as the standardizer:

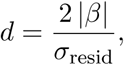

where *β* is the standardized regression coefficient with both predictor and outcome *z*-scored before model fitting and *σ*_resid_ is the square root of the mean squared error from the covariate-adjusted OLS model. Values of *d* are reported as magnitudes; the direction of each effect is given by the sign of *β*.

The lead author used OpenAI Codex (GPT-6.1 Sol) and Devin (Cognition AI; Adaptive mode with automatic model selection) to assist with reviewing and revising data-processing and analysis scripts. Codex also assisted with refining plotting scripts. The authors determined the final analytical procedures; the lead author reviewed and edited the AI-assisted output and verified the code, results and figures.

### 2.11 Apparatus

The task was programmed using PsychoPy v2023.1^23^ and deployed online through Pavlovia (Open Science Tools Ltd., Nottingham, UK). Participants completed the study remotely in a web browser. They were instructed to use a desktop or laptop computer, complete the task in fullscreen mode, and respond using a physical keyboard. The task was optimized for desktop browser use, and timing was monitored using PsychoJS timing routines and frame checks.

### 2.12 Ethics

The study was approved by the Stanford University Institutional Review Board (Protocol #70039; initially approved May 26, 2023, with subsequent amendments). All participants provided informed consent electronically. Participants were compensated via Prolific at $15 per hour.

## 3 Results

### 3.1 Participant characteristics

The final analytic sample included 497 participants (Table 1). Participants were 22.0 *±* 1.7 yrs old (mean *±* SD), the sample was 49.7% female (*N* = 247) and 50.3% male (*N* = 250), and mean education was 14.8 *±* 2.1 yrs (*N* = 496). Table 1 presents demographic and usage descriptives by aggregate-use group.

**Table 1:**
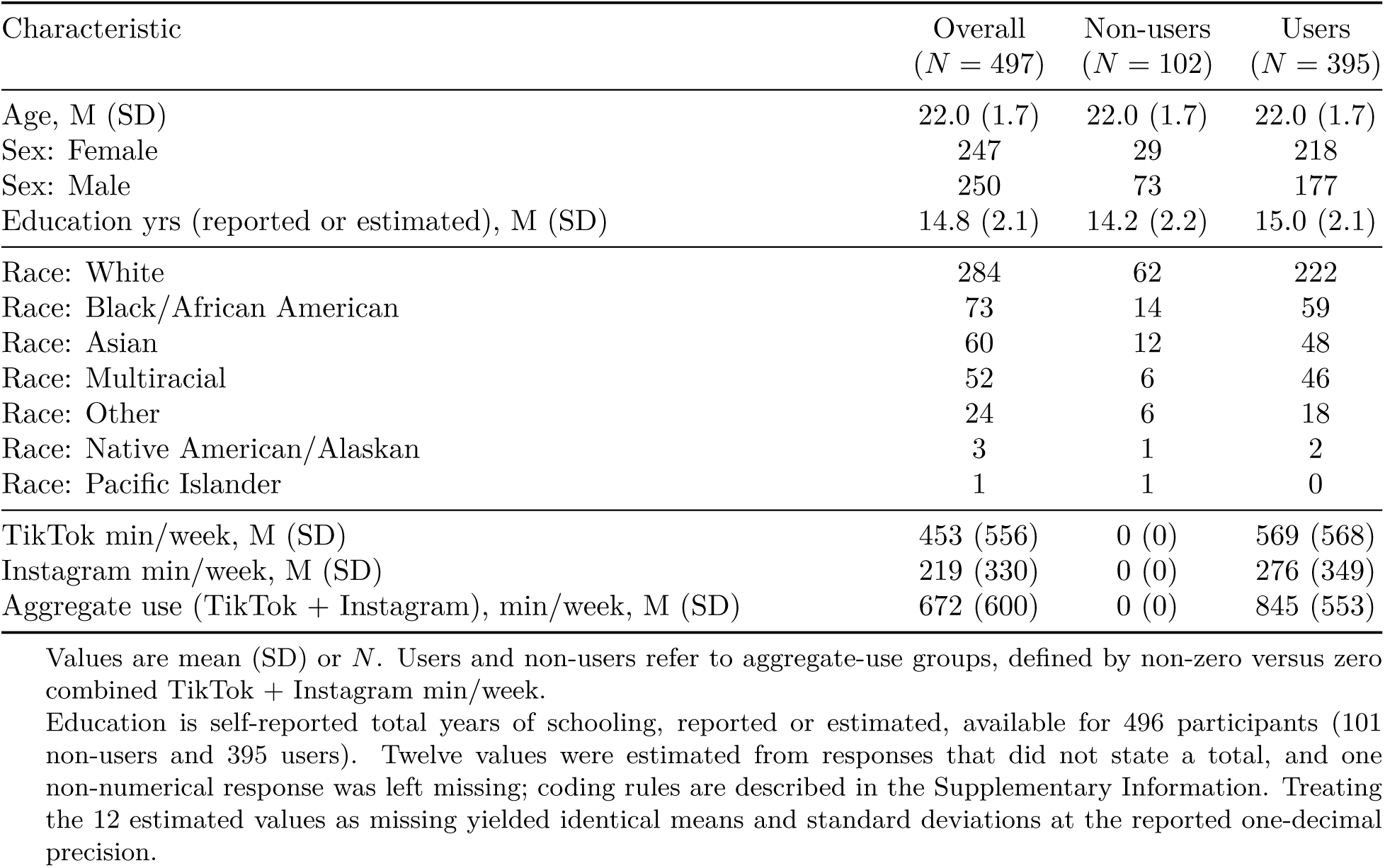
Participant characteristics and descriptive statistics.

| Characteristic | Overall<br>(*N* = 497) | Non-users<br>(*N* = 102) | Users<br>(*N* = 395) |
|---|---|---|---|
| Age, M (SD) | 22.0 (1.7) | 22.0 (1.7) | 22.0 (1.7) |
| Sex: Female | 247 | 29 | 218 |
| Sex: Male | 250 | 73 | 177 |
| Education yrs (reported or estimated), M (SD) | 14.8 (2.1) | 14.2 (2.2) | 15.0 (2.1) |
| Race: White | 284 | 62 | 222 |
| Race: Black/African American | 73 | 14 | 59 |
| Race: Asian | 60 | 12 | 48 |
| Race: Multiracial | 52 | 6 | 46 |
| Race: Other | 24 | 6 | 18 |
| Race: Native American/Alaskan | 3 | 1 | 2 |
| Race: Pacific Islander | 1 | 1 | 0 |
| TikTok min/week, M (SD) | 453 (556) | 0 (0) | 569 (568) |
| Instagram min/week, M (SD) | 219 (330) | 0 (0) | 276 (349) |
| Aggregate use (TikTok + Instagram), min/week, M (SD) | 672 (600) | 0 (0) | 845 (553) |

Values are mean (SD) or *N*. Users and non-users refer to aggregate-use groups, defined by non-zero versus zero combined TikTok + Instagram min/week.
Education is self-reported total years of schooling, reported or estimated, available for 496 participants (101 non-users and 395 users). Twelve values were estimated from responses that did not state a total, and one non-numerical response was left missing; coding rules are described in the Supplementary Information. Treating the 12 estimated values as missing yielded identical means and standard deviations at the reported one-decimal precision.

Short-form platform engagement was widespread: 58.1% were classified as TikTok users (*N* = 289), 58.6% as Instagram Reels users (*N* = 291), 53.5% as YouTube Shorts users (*N* = 266) and 14.1% as Facebook Reels users (*N* = 70). Most participants were classified as users of two or more short-form video platforms (*N* = 303; 61.0%; Fig. 3**A**). Of 497 participants, 395 (79.5%) were device-reported “aggregate-use users” (non-zero TikTok + Instagram minutes) and 102 (20.5%) were “aggregate-use non-users” (zero use on both platforms). Total aggregate use averaged 845 *±* 553 min/week among aggregate-use users (Fig. 3**B**). The 395 users were divided into tertiles based on aggregate use: Low (*N* = 132; 2–518 min/week), Medium (*N* = 131; 519–1,022 min/week) and High (*N* = 132; 1,026–2,550 min/week) (Fig. 3**C**). Device-reported and self-reported use were modestly correlated for TikTok (*r* = 0.31, *p <* 0.001, *N* = 286); device-reported Instagram use was likewise modestly correlated with self-reported Instagram Reels use (*r* = 0.30, *p <* 0.001, *N* = 290), suggesting that retrospective self-report captured a limited proportion of variance in device-reported screen time (Fig. 3**D**).

**Figure 3:**
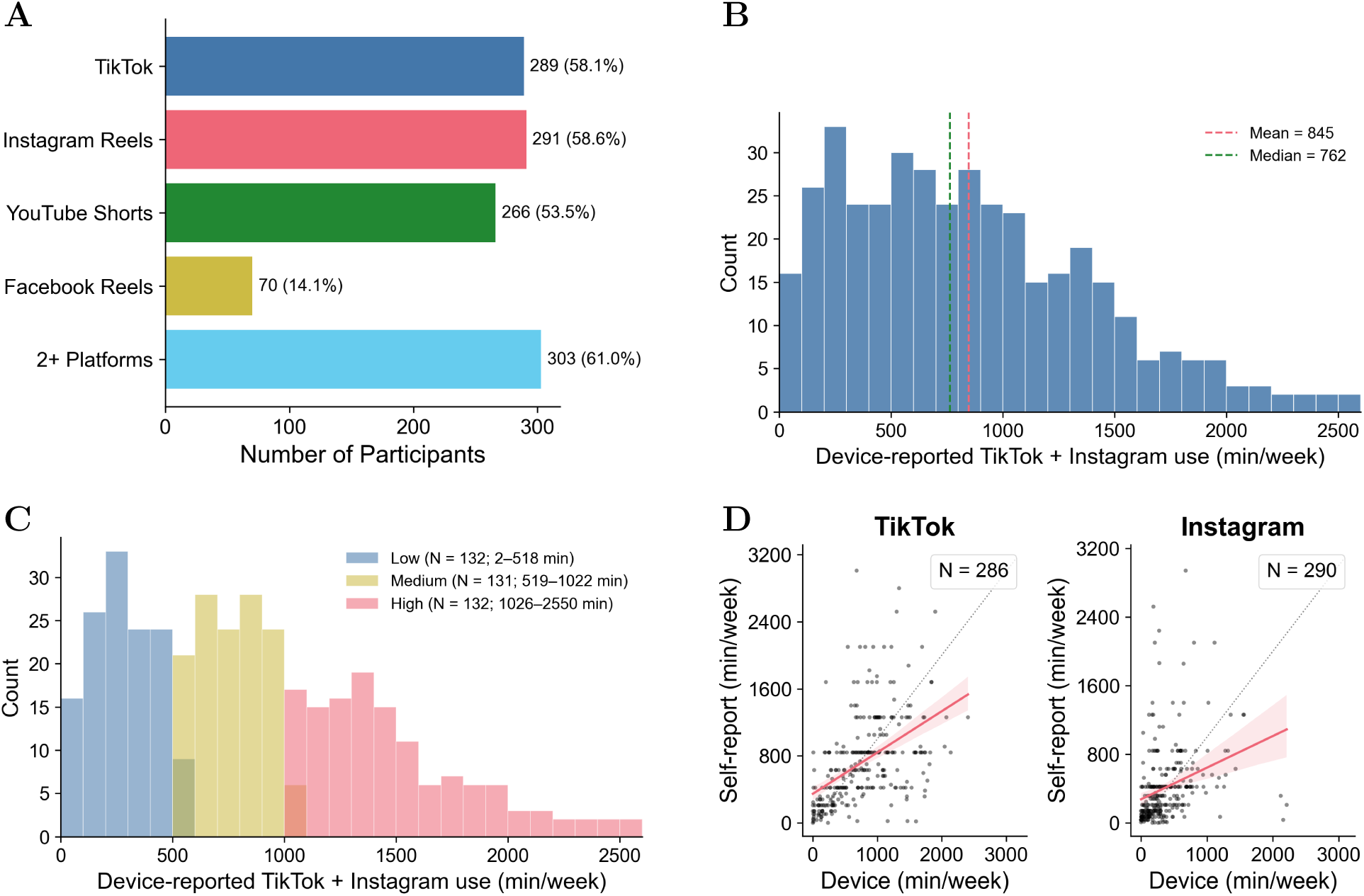
Short-form video use in the Prolific sample. **A**, Non-exclusive classified platform usage across the final analytic sample, showing the number of participants (*N* = 497) classified as users of TikTok, Instagram, YouTube Shorts and Facebook Reels. Participants self-reporting TikTok or Instagram use but recording zero device-reported minutes were classified as non-users of that platform. **B**, Distribution of aggregate short-form video use, defined as TikTok plus Instagram minutes per week, among users (*N* = 395; mean = 845 min/week, median = 762 min/week, SD = 553 min/week). **C**, The same aggregate-use distribution shown in **B**, colored by usage tertile. Panels **B** and **C** use identical bin edges and x-axis limits. **D**, Comparison of device-reported and retrospective self-reported screen time for TikTok, and of device-reported Instagram screen time with retrospective self-reported Instagram Reels use. The two highest self-reported TikTok values (8,400 and 5,460 min/week) were excluded from the visualization (*N* = 284 shown; *N* = 286 analyzed); all Instagram observations are shown (*N* = 290).

### 3.2 Short-form video use and sustained attention

To investigate the relationship between sustained attention and short-form video use, we first conducted participant-level linear regressions among aggregate-use users, with sex included as a covariate (Fig. 4). Device-reported aggregate short-form video use was not significantly associated with attention *d^′^*(*β* = *−*0.006, 95% CI [*−*0.108, 0.097], *p* = 0.92), frequent-trial attention accuracy (*β* = *−*0.045, 95% CI [*−*0.147, 0.057], *p* = 0.38) or infrequent-trial attention accuracy (*β* = 0.006, 95% CI [*−*0.096, 0.108], *p* = 0.90). We note that the performance range was restricted for frequent-trial accuracy. Sensitivity analysis indicated 80% power to detect an absolute partial correlation of *r* = 0.14 for the usage–attention *d^′^*association; smaller associations may have gone undetected.

**Figure 4:**
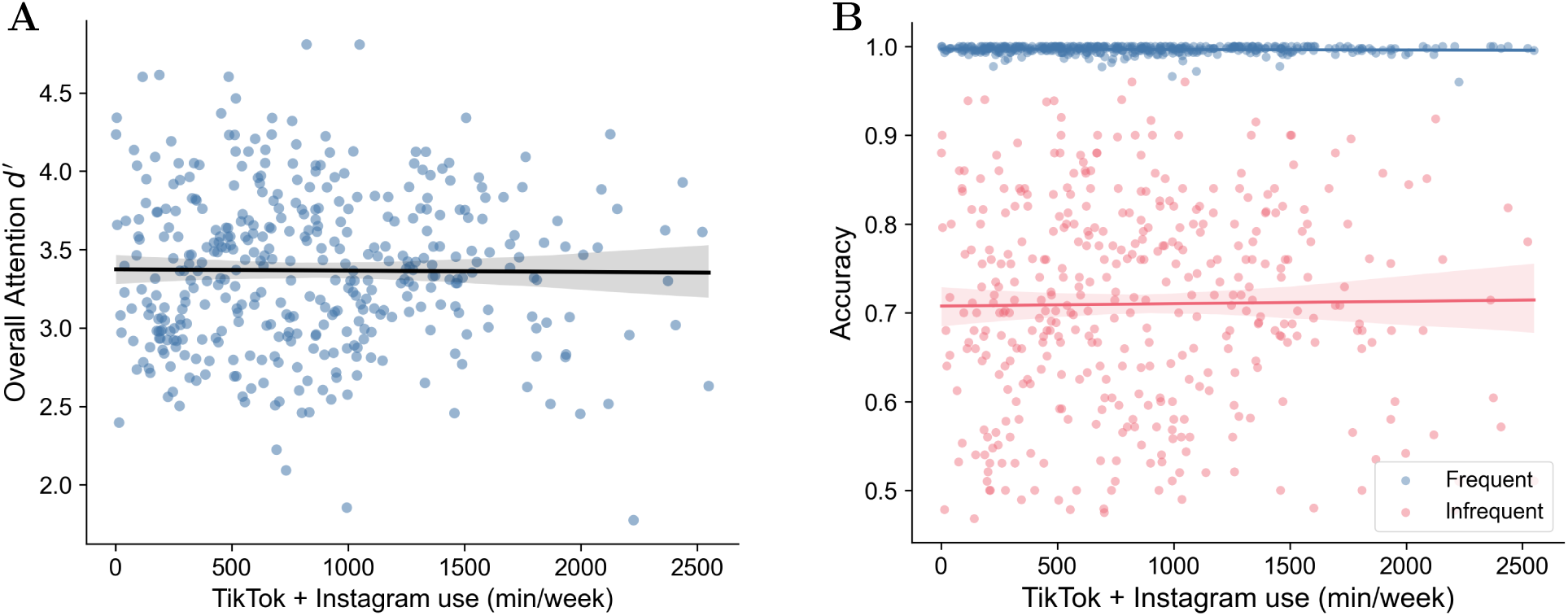
Sustained-attention outcomes as a function of short-form video use. **A**, Attention *d^′^* versus aggregate short-form video use among aggregate-use users (*β* = *−*0.01, *p* = 0.92, *N* = 395). **B**, Frequent-and infrequent-trial attention accuracy versus aggregate short-form video use among aggregate-use users. Frequent accuracy: *β* = *−*0.05, *p* = 0.38; infrequent accuracy: *β* = 0.01, *p* = 0.90. Points are participant observations; lines are unadjusted ordinary least-squares visualization fits and shaded bands are their 95% bootstrap confidence intervals. Reported coefficients and *p* values are from the corresponding sex-adjusted models.

To examine sustained attention at the trial level, we next conducted a linear mixed-effects model among aggregate-use users (*N* = 395 participants; 197,500 trial observations). Binary trial-level attention accuracy (correct = 1, incorrect = 0) was modeled as a function of trial condition (frequent vs. infrequent), continuous aggregate use, sex, and the condition *×* usage and condition *×* sex interactions, with a random intercept for participant. As expected, accuracy was lower for infrequent than frequent trials. Across the modeled trials (all 500 trials, with no-response trials scored as incorrect), participant-level mean accuracy was 0.980 (SD = 0.018) for frequent trials and 0.687 (SD = 0.122) for infrequent trials. Notably, aggregate use was not significantly associated with attention accuracy (*b* = *−*0.0012 per 1,000 min/week, SE = 0.0022, 95% CI [*−*0.0055, 0.0030], *p* = 0.57), and the condition *×* usage interaction was not significant (*b* = 0.0004 per 1,000 min/week, SE = 0.0027, 95% CI [*−*0.0050, 0.0058], *p* = 0.89).

Finally, secondary full-sample analyses also revealed no significant differences between aggregate-use users and non-users in attention *d^′^*(*b* = *−*0.022, 95% CI [*−*0.134, 0.089], *p* = 0.697), frequent-trial attention accuracy (*b* = 0.0004, 95% CI [*−*0.0006, 0.0014], *p* = 0.438), or infrequent-trial attention accuracy (*b* = *−*0.019, 95% CI [*−*0.045, 0.007], *p* = 0.155). These categorical comparisons were treated as secondary because aggregate-use non-users were defined by zero TikTok and Instagram use rather than zero use across all short-form video platforms.

Collectively, the preceding analyses did not reveal a significant relationship between aggregate use and sustained-attention performance in this sample. To further examine this null relationship, a Bayesian multiple linear regression was conducted among aggregate-use users (*N* = 395) to estimate the relationship between aggregate use and attention *d^′^*, controlling for sex, using the brms package^21^ in R. Continuous variables were standardized prior to analysis to yield standardized posterior estimates. The standardized posterior estimate was close to zero (*β* = *−*0.006, 95% CrI [*−*0.109, 0.099]). Exploratory analyses likewise found no significant associations between aggregate use and median reaction time or reaction-time variability, either overall or within the frequent and infrequent conditions (Supplementary Figs. S2 and S3).

### 3.3 Short-form video use is associated with poorer episodic memory

To investigate the relationship between episodic memory and short-form video use, we first conducted participant-level linear regressions among aggregate-use users, with sex included as a covariate. Greater device-reported aggregate short-form video use was negatively associated with overall memory *d^′^*(*β* = *−*0.150, *p* = 0.004, 95% CI [*−*0.251, *−*0.049], Cohen’s *d* = 0.303), indicating that heavier users demonstrated poorer recognition sensitivity (Fig. 5**A**).

**Figure 5:**
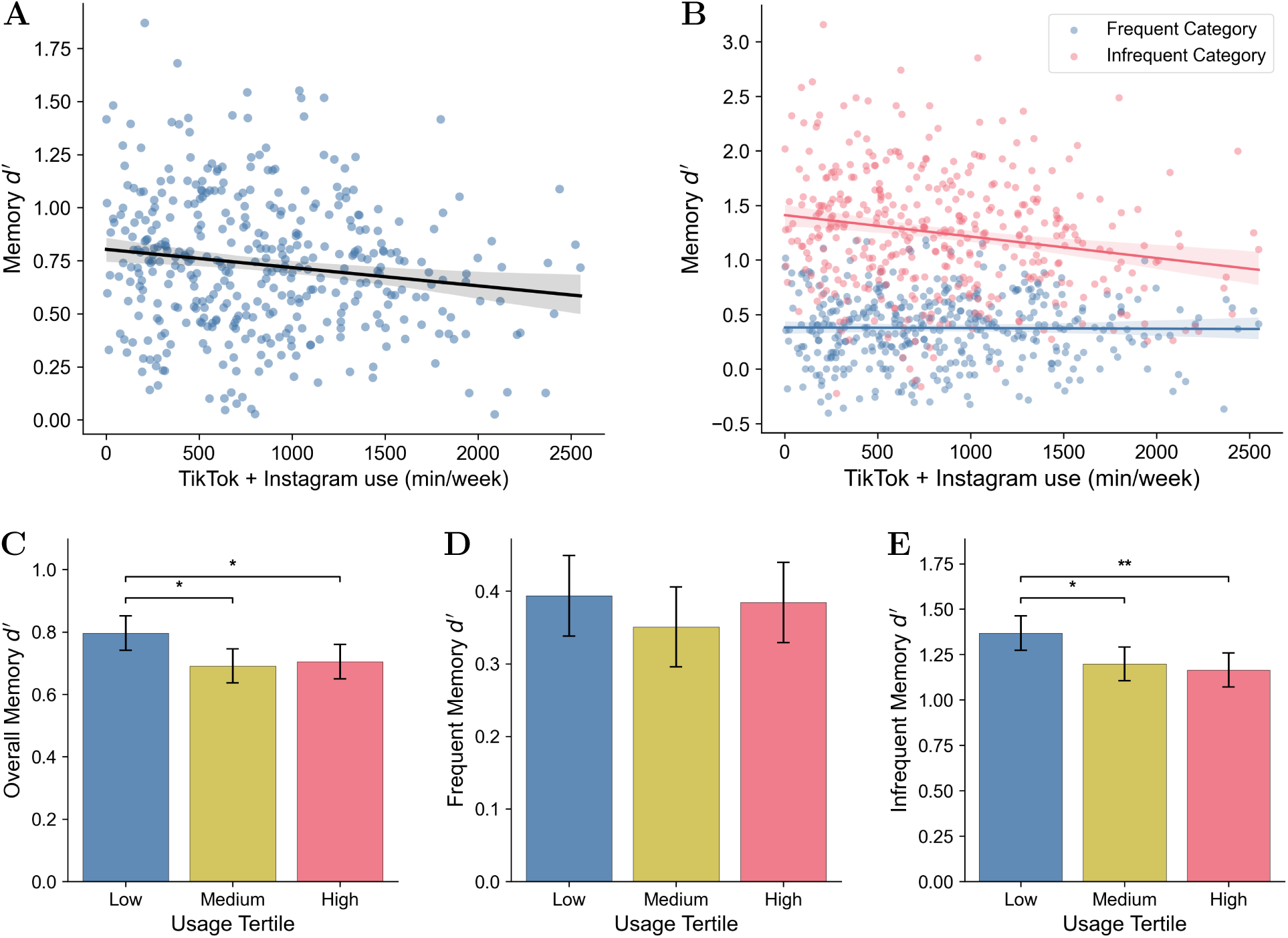
Greater aggregate short-form video use is associated with poorer episodic memory. **A**, Overall memory *d^′^* versus aggregate short-form video use among aggregate-use users (*β* = *−*0.15, *p* = 0.004, 95% CI [*−*0.25, *−*0.05], *N* = 395). **B**, Frequent-category and infrequent-category memory *d^′^* versus aggregate short-form video use. Frequent-category memory: *β* = *−*0.04, *p* = 0.41; infrequent-category memory: *β* = *−*0.18, *p <* 0.001, 95% CI [*−*0.28, *−*0.08]. In **A–B**, points are participant observations; lines are unadjusted ordinary least-squares visualization fits and shaded bands are their 95% bootstrap confidence intervals, whereas reported coefficients and *p* values are from sex-adjusted models. **C–E**, Sex-adjusted estimated marginal means with model-based 95% confidence intervals for overall (**C**), frequent-category (**D**) and infrequent-category (**E**) memory *d^′^* by aggregate-use tertile. Significance brackets denote Holm-corrected pairwise comparisons from the sex-adjusted OLS models: * *p*_Holm_ *<* 0.05 and ** *p*_Holm_ *<* 0.01.

To test whether these participant-level associations differed by memory condition (Fig. 5**B**), we fit a repeated-measures model of condition-specific memory *d^′^*. The sex-adjusted condition *×* usage interaction was significant (*b* = *−*0.086, SE = 0.028, 95% CI [*−*0.142, *−*0.031], *p* = 0.002), indicating that the negative association between aggregate use and memory was stronger for infrequent-category than frequent-category items. To characterize this condition-specific pattern, we then conducted separate OLS regressions for frequent-and infrequent-category memory *d^′^*. Greater aggregate use was negatively associated with infrequent-category (outdoor) memory *d^′^* (*β* = *−*0.182, *p <* 0.001, 95% CI [*−*0.282, *−*0.082], Cohen’s *d* = 0.371). In contrast, frequent-category memory *d^′^* showed no significant association with use (*β* = *−*0.043, *p* = 0.408). These findings raise the possibility that greater aggregate use is differentially associated with lower memory performance for infrequent-category items, though we note that interpretative caution is warranted here given that the low level of performance for frequent-category items creates a restricted range.

To examine recognition accuracy across frequent-and infrequent-category items, we next conducted a trial-level linear mixed-effects model of recognition accuracy among aggregate-use users (*N* = 395 participants; 79,000 trial observations), with condition, aggregate use, sex, and the condition *×* use and condition *×* sex interactions as fixed effects and participant as a random intercept. In general, recognition accuracy was higher for infrequent-category than frequent-category items: participant-level mean recognition accuracy was 0.565 (SD = 0.057) for frequent-category items and 0.694 (SD = 0.085) for infrequent-category items. The condition *×* usage interaction was significant (unstandardized accuracy-scale coefficient *b* = *−*0.022 per 1,000 min/week, SE = 0.006, 95% CI [*−*0.034, *−*0.010], *p <* 0.001). Within the frequent category, aggregate use was not significantly associated with recognition accuracy (*b* = *−*0.004 per 1,000 min/week, SE = 0.006, 95% CI [*−*0.016, 0.008], *p* = 0.56); within the infrequent category, aggregate use was significantly negatively associated with recognition accuracy (*b* = *−*0.026 per 1,000 min/week, SE = 0.006, 95% CI [*−*0.038, *−*0.013], *p <* 0.001). As such, the association between aggregate use and recognition accuracy was more negative for infrequent-category than frequent-category items.

Next, we examined group differences in memory performance among low, medium and high users. For overall memory *d^′^*, mean performance was 0.80 in the Low group, 0.69 in the Medium group and 0.70 in the High group (Fig. 5**C**). A sex-adjusted omnibus model showed a significant effect of usage tertile, *F* (2, 391) = 4.04, *p* = 0.018. Holm-corrected pairwise comparisons indicated that the Low usage group had better memory performance than both the Medium usage group (*p*_Holm_ = 0.026) and the High usage group (*p*_Holm_ = 0.048), whereas the Medium and High groups did not differ (*p*_Holm_ = 0.727). Frequent-category memory *d^′^* did not significantly differ across tertiles, *F* (2, 391) = 0.66, *p* = 0.518 (Low = 0.38, Medium = 0.35, High = 0.39; Fig. 5**D**). In contrast, infrequent-category memory *d^′^* differed significantly across tertiles, *F* (2, 391) = 5.07, *p* = 0.007 (Low = 1.38, Medium = 1.20, High = 1.16; Fig. 5**E**).

Secondary full-sample analyses found no significant differences between aggregate-use users and non-users in overall memory *d^′^* (*b* = 0.000, 95% CI [*−*0.072, 0.073], *p* = 0.996), frequent-category memory *d^′^* (*b* = 0.028, 95% CI [*−*0.045, 0.101], *p* = 0.450), or infrequent-category memory *d^′^* (*b* = *−*0.049, 95% CI [*−*0.173, 0.075], *p* = 0.438). Among aggregate-use users, mean overall, frequent-category and infrequent-category memory *d^′^* values were 0.731 (SD = 0.321), 0.376 (SD = 0.322) and 1.244 (SD = 0.550), respectively. Among non-users, the corresponding values were 0.731 (SD = 0.338), 0.322 (SD = 0.363) and 1.317 (SD = 0.576).

### 3.4 Short-form video use predicts memory after accounting for sustained-attention performance

We next investigated whether the association between aggregate use and memory remained after controlling for sustained-attention performance.

In a linear model with overall memory *d^′^* as the outcome measure, both overall attention *d^′^* (*β* = 0.251, *p <* 0.001, 95% CI [0.156, 0.346], Cohen’s *d* = 0.523) and aggregate use (*β* = *−*0.149, *p* = 0.003, 95% CI [*−*0.247, *−*0.051], Cohen’s *d* = 0.310) were significant predictors of memory performance after controlling for sex.

To test whether the condition-specific usage–memory association persisted after accounting for sustained attention, we fit a participant-level repeated-measures model that included overall attention *d^′^* and allowed attention to interact with memory condition. The condition *×* usage interaction remained significant (*b* = *−*0.086, SE = 0.028, 95% CI [*−*0.140, *−*0.032], *p* = 0.002). For infrequent-category memory *d^′^*, both infrequent-trial attention accuracy (*β* = 0.271, *p <* 0.001, 95% CI [0.177, 0.364], Cohen’s *d* = 0.573) and aggregate use (*β* = *−*0.184, *p <* 0.001, 95% CI [*−*0.280, *−*0.087], Cohen’s *d* = 0.389) were significant predictors after controlling for sex. By contrast, for frequent-category memory *d^′^*, frequent-trial attention accuracy was not a significant predictor (*β* = 0.034, *p* = 0.498, 95% CI [*−*0.065, 0.133]), and aggregate use was not significant (*β* = *−*0.041, *p* = 0.426, 95% CI [*−*0.143, 0.060]). These results indicate that the stronger negative association between aggregate use and memory for infrequent-category relative to frequent-category items remained after accounting for overall sustained-attention performance.

### 3.5 Self-reported short-form video use and cognition

As device-reported and self-reported usage were only modestly correlated, we repeated the primary participant-level analyses using self-reported combined TikTok and Instagram Reels use (TikTok + Instagram Reels minutes per week). Among self-reported aggregate-use users (*N* = 399; *M* = 866 min/week, SD = 794), self-reported aggregate use was not associated with attention *d^′^* (*β* = *−*0.033, *p* = 0.516), frequent-trial attention accuracy (*β* = *−*0.093, *p* = 0.066), or infrequent-trial attention accuracy (*β* = 0.002, *p* = 0.970). For episodic memory, self-reported aggregate use was not significantly associated with overall memory *d^′^* (*β* = *−*0.057, *p* = 0.260), frequent-category memory *d^′^* (*β* = *−*0.003, *p* = 0.960), or infrequent-category memory *d^′^*(*β* = *−*0.090, *p* = 0.074). A trial-level linear mixed-effects model among these participants (*N* = 399; 79,800 trial observations) yielded a condition *×* self-reported aggregate use interaction that did not reach significance (*b* = *−*0.0084 per 1,000 min/week, SE = 0.0043, 95% CI [*−*0.0167, 0.0000], *p* = 0.051). Linear contrasts from this same interaction model showed no usage association for frequent-category accuracy (*b* = 0.0006, SE = 0.0042, 95% CI [*−*0.0077, 0.0089], *p* = 0.886) and a nonsignificant negative association for infrequent-category accuracy (*b* = *−*0.0077, SE = 0.0042, 95% CI [*−*0.0161, 0.0006], *p* = 0.068). Parallel analyses among participants reporting nonzero self-reported TikTok use (*N* = 301) showed the same qualitative pattern but with weaker effects (attention *d^′^*: *β* = *−*0.022, *p* = 0.703; infrequent-category memory *d^′^*: *β* = *−*0.080, *p* = 0.170). In exploratory sensitivity analyses using device-reported TikTok minutes alone, associations were weaker than for the combined measure and did not reach significance: among TikTok users (*N* = 289), TikTok use was not significantly associated with infrequent-category memory *d^′^*(*β* = *−*0.109, *p* = 0.071), overall memory *d^′^*(*β* = *−*0.071, *p* = 0.246) or any attention outcome (Supplementary Table S4).

### 3.6 Addiction-like engagement, media multitasking and impulsivity

Finally, we examined whether TikTok addiction-like engagement (TikTok Addiction Scale), media multitasking and/or impulsivity were associated with aggregate use. TikTok Addiction Scale (TAS) scores were positively associated with TikTok minutes among TikTok users (*β* = 0.38, *p <* 0.001, 95% CI [0.27, 0.49], Cohen’s *d* = 0.85, *N* = 289; Fig. 6**A**). Adapted MMI scores showed a weak positive association with aggregate use (*β* = 0.13, *p* = 0.014, 95% CI [0.026, 0.229], Cohen’s *d* = 0.26, *N* = 395; Fig. 6**B**).

**Figure 6:**
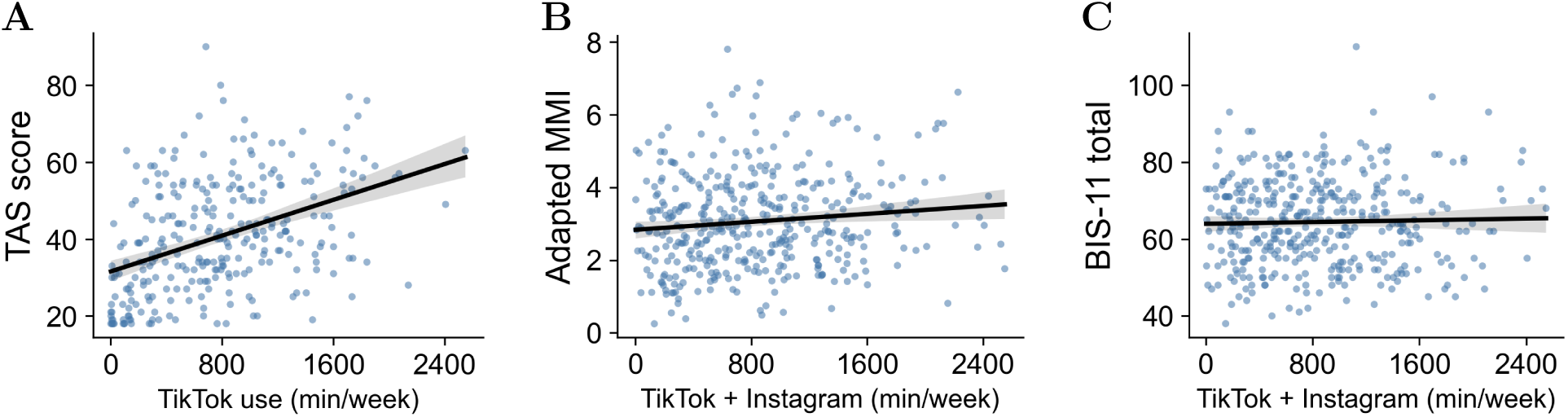
Addiction-like engagement, media multitasking and impulsivity measures. **A**, TikTok Addiction Scale (TAS) scores versus TikTok minutes among TikTok users included in the TAS analysis (*β* = 0.38, *p <* 0.001, 95% CI [0.27, 0.49], Cohen’s *d* = 0.85, *N* = 289). **B**, Adapted Media Multitasking Index (MMI) scores versus aggregate short-form video use (*β* = 0.13, *p* = 0.014, 95% CI [0.026, 0.229], Cohen’s *d* = 0.26, *N* = 395). **C**, BIS-11 impulsivity scores versus aggregate short-form video use (*β* = 0.04, *p* = 0.486, *N* = 395). Points are participant observations; lines are unadjusted ordinary least-squares visualization fits and shaded bands are their 95% bootstrap confidence intervals. Reported coefficients and *p* values are from sex-adjusted models.

Given past observations that episodic memory is lower in heavier media multitaskers ^20,24^, we tested whether the observed relationships between device-reported usage and memory held after controlling for media multitasking by fitting models predicting overall and infrequent-category memory *d^′^* from both aggregate use and adapted MMI scores, with sex as a covariate. Aggregate use remained a significant predictor of overall memory *d^′^* (*β* = *−*0.15, *p* = 0.004) and infrequent-category memory *d^′^* (*β* = *−*0.18, *p <* 0.001), whereas MMI score was not a significant predictor in either model (overall *d^′^*: *β* = *−*0.005, *p* = 0.924; infrequent *d^′^*: *β* = *−*0.02, *p* = 0.700). These results indicate that the association between aggregate use and memory remained significant after controlling for adapted MMI scores. BIS-11 impulsivity scores were not significantly associated with aggregate use (*β* = 0.04, *p* = 0.486, *N* = 395; Fig. 6**C**). All models included sex as a covariate.

## 4 Discussion

Driven by the global ubiquity of platforms such as TikTok, Instagram and YouTube, short-form video formats have emerged as a defining element of modern communication, entertainment and social interaction. In the present study, 79.5% of participants had device-reported nonzero aggregate TikTok or Instagram use, and aggregate use across the full sample (including aggregate-use non-users) averaged 672 min/week. These findings align with recent demographic data indicating widespread engagement with social media platforms among young adults ^3^. Although short-form video has been broadly adopted, empirical studies offer conflicting conclusions about whether intensive use is associated with poorer cognitive function ^1^.

We investigated whether individual differences in short-form video use relate to sustained attention and episodic memory performance. We report that greater device-reported aggregate short-form video use was associated with poorer episodic memory performance, with the strongest association for infrequent-category items. Conversely, device-reported aggregate short-form video use did not significantly relate to sustained attention metrics. An earlier locally collected college-age pilot study showed a broadly similar pattern: greater TikTok use was associated with lower overall and infrequent-category memory *d^′^*, whereas attention *d^′^* was not significantly associated with use (Supplementary Table S2).

This selectivity is consistent with an attention–memory coupling account, but it may also reflect the novelty and distinctiveness of infrequent-category items. On the attention–memory coupling interpretation, lapses or reductions in controlled attention would be expected to disproportionately impair memory for infrequent-category items, because those items require stronger attentional engagement and response override than the frequent-category items that support prepotent responding. In the SART-M, infrequent-category items were both relatively rare and encountered under conditions requiring greater attentional engagement and cognitive control (e.g., to overcome the prepotent categorization and response tendency); the present design therefore cannot isolate novelty-related encoding from attentional demand. Because outdoor scenes were always the infrequent category, scene-category encoding differences also cannot be ruled out. Greater short-form video use may be associated with a reduced mnemonic benefit from novelty, greater vulnerability under demanding encoding conditions, or both.

One possibility is that heavier short-form video use is associated with greater exposure to large volumes of unrelated and rapidly changing information. Such fragmented information processing could increase interference or reduce distinctive processing, potentially attenuating the mnemonic benefit of novelty. Novelty detection is thought to promote episodic encoding by signaling that new information should be encoded into memory ^25^. If heavy short-form video use were to reduce novelty responsivity, the mnemonic benefit typically associated with distinctive events could be diminished. This possibility is consistent with work showing an attenuated P300 oddball response in heavy short-form-video users relative to lower-frequency users ^6^, although that study framed its findings in terms of attention-related processing rather than episodic novelty encoding. Future work will need to directly distinguish the contributions of attentional demand, novelty detection and information overload to memory performance in short-form video users.

In contrast to the observed memory effects, sustained-attention outcomes did not show reliable associations with device-reported aggregate short-form video use. This null pattern for attention, combined with the reliable memory associations, raises the possibility that the pattern associated with short-form video use may be more evident in downstream encoding outcomes than in sustained-attention performance. One interpretation is that stable individual differences in baseline sustained-attention capacity may outweigh any variation associated with short-form video use on task performance. Alternatively, cumulative encoding consequences during the SART-M, captured by the surprise memory test, may amplify subtle within-task fluctuations into measurable between-person variability. This interpretation is aligned with prior work linking sustained attention to subsequent or incidental memory performance ^13,26^. An exploratory post hoc analysis adapting the participant-level approach of deBettencourt et al. was not significant (Supplementary Fig. S5).

Critically, the relationship between infrequent-category memory performance and device-reported aggregate short-form video use remained significant after accounting for sustained-attention performance. Overall attention *d^′^* predicted overall memory *d^′^*, and infrequent-trial attention accuracy predicted infrequent-category memory *d^′^*. Frequent-trial attention accuracy (which was at ceiling) did not significantly predict frequent-category memory *d^′^* (which was near floor). However, device-reported aggregate short-form video use continued to negatively predict overall and infrequent-category memory *d^′^* after controlling for attention and sex. This pattern suggests that heavier short-form video use was not simply a proxy for poorer sustained attention in the task; rather, it accounted for additional variance in memory performance, especially for infrequent-category items. The association also remained after adjustment for adapted media-multitasking scores.

Device-reported aggregate short-form video use was not significantly associated with sustained-attention performance as measured by the SART-M. For attention *d^′^*, the 95% confidence interval ranged from *−*0.108 to 0.097; the model had 80% power to detect *|r|* = 0.14, so smaller associations cannot be ruled out. The SART-M indexes only one facet of attention. Prior behavioral work in this area examined several distinct attentional constructs rather than sustained attention alone. Thomson et al., ^8^ using both self-report and objective smartphone usage measures, found no evidence that social media engagement predicted attentional bias toward social media stimuli. Similarly, Jones et al., ^9^ using objective mobile-use measures, found only small associations between social media use and attentional control, with greater overall social media/TikTok use associated with slightly better, rather than worse, attentional control. In contrast, other short-form video studies report poorer self-reported attentional control ^4^, deficits during Stroop and video-watching tasks ^5^, attenuated P300 responses during a visual oddball task ^6^, and poorer sustained-attention performance in the survey phase of Lin et al.,^7^ including increased commission errors and reaction-time variability on a sustained attention to response task. This complex mix of findings raises the possibility that short-form digital media consumption may be selectively related to particular facets of attention rather than to a generalized sustained-attention deficit.

In the present study, comparisons of device-reported aggregate short-form video users and non-users revealed no significant differences in attention or memory performance, whereas continuous variation in short-form video use among users was associated with memory performance. This pattern suggests that a binary distinction between users and non-users may be insufficient to characterize the observed association. However, because medium-and high-use groups did not differ from one another, the present findings should not be interpreted as evidence for a simple monotonic dose–response relationship between short-form video use and cognition. Longitudinal work in adolescents has similarly identified associations between increasing social-media-use trajectories and performance in several cognitive domains^27^, although such observational findings do not establish causality.

To assay short-form video use, we collected both retrospective self-report measures and device-reported screenshot-based measures. The two measures were only modestly correlated, indicating that retrospective self-report captured limited variance in device-reported usage. This finding underscores a critical methodological need for objective or device-verified measures of social media use, rather than relying primarily on retrospective self-report ^9^. In the current study, we examined device-reported aggregate short-form video use across TikTok and Instagram because both platforms prominently feature algorithmically curated short-form video^1^. However, other platforms, including YouTube and Facebook, increasingly feature short-form content alongside traditional media formats. As a result, our aggregate TikTok and Instagram measure does not capture participants’ total cross-platform exposure to short-form video content. Future studies should distinguish short-form video viewing from other activity within each application and capture total exposure across platforms.

Notably, most of the device-reported “non-users” of TikTok and Instagram in the present study nonetheless self-reported use of at least one other short-form video platform, including YouTube Shorts and Facebook Reels, raising the possibility that the null main effect of users vs. “non-users” on episodic memory reflects engagement with short-form video on other platforms. Restricting non-users to participants reporting zero use across all four named short-form video platforms reduced the group to *N* = 32, reflecting substantial use of other short-form video platforms among aggregate-use non-users.

Platform-level algorithmic differences are also important to consider. Although TikTok, Instagram (via Reels), YouTube (via Shorts) and Facebook (via Reels) all include short-form video content, they differ in recommendation systems, content ecosystems, social affordances and the extent to which short-form video is central to the user experience. TikTok provides a relatively dedicated short-form video feed, whereas Instagram, YouTube and Facebook combine short-form videos with other content formats (e.g., Stories, longer-form video, social-network feeds). These differences make it difficult to assume that a minute of use has the same cognitive or experiential meaning across platforms. For this reason, the primary analyses focused on TikTok and Instagram, for which we collected app-or device-based screen-time measures; however, broader cross-platform exposure should be examined more directly in future work. TikTok-only and platform-specific analyses showed weaker associations overall (Supplementary Tables S4 and S5); differences in statistical significance do not establish that an aggregate measure is superior.

One possible mechanism that could contribute to the observed usage–memory association is that rapid, fast-paced short-form video viewing can involve frequent context switching, which may disrupt episodic encoding and memory integration ^28,29^. For instance, short-form video viewing has been shown to be negatively related to prospective memory, suggesting that rapidly switching contexts may interfere with intention maintenance and later recall ^10,28^. Relatedly, participants who viewed multiple short videos matched in duration and content later showed poorer recall than participants who viewed a continuous long video, alongside reduced retrieval-related neural activity and connectivity in regions implicated in information integration and cognitive control ^29^. In the current study, the rapidly switching stimuli presented during the sustained-attention task may partially mirror the fragmented structure of short-form video. It is therefore possible that the SART-M was particularly sensitive to memory-encoding differences in high-use participants, given its rapidly changing stimuli. This possibility was not tested in the present study.

Our findings indicate that the relationship between aggregate short-form video use and episodic memory is small, highlighting the necessity of large-scale datasets to robustly capture individual differences. Furthermore, the direction of causality remains unclear: short-form video engagement may alter memory processes over time, individuals with baseline differences in memory may be drawn to short-form video, or both pathways may contribute to the relationship. Longitudinal studies of adolescents have found associations between digital media use and later ADHD symptoms ^30^, as well as associations between habitual social media checking and longitudinal functional brain development in salience, motivation and cognitive control networks ^31^. Both use and memory could also reflect unmeasured third variables, such as sleep, overall screen time, ADHD symptoms or engagement with the online task.

Neuroimaging studies have also identified neural responses associated with short-form video and broader social media engagement. For example, viewing personalized relative to non-personalized TikTok clips was associated with greater activation in default-mode network subregions and the ventral tegmental area ^32^, whereas a small naturalistic study found reduced executive-function accuracy and altered prefrontal activation following brief social media exposure ^33^. Together with emerging evidence of neuroanatomical and functional correlates of short-form video addiction^34^, these findings raise the possibility that sustained engagement with short-form video may be associated with longer-term neural and cognitive differences. The present study did not measure neural function or long-term cognitive change.

Several limitations should be noted. First, in the current study, aggregate-use non-users were defined as participants with device-reported 0 minutes of TikTok and Instagram use, rather than 0 minutes across all short-form video platforms. Most participants in this group self-reported use of at least one other short-form video platform, such as YouTube Shorts or Facebook Reels. Device-based use was not collected for YouTube, Facebook or other platforms, and the Instagram measure includes activity beyond Reels. Accordingly, the primary analyses emphasized continuous variation in TikTok and Instagram use among aggregate-use users rather than categorical user/non-user contrasts. Second, usage values were entered by participants from screen-time reports and checked against uploaded screenshots. These reports did not uniformly cover seven complete days; some weekly values were estimated from daily averages. A single week may not represent habitual use, and screen-time reports do not capture how attentively content was viewed. Third, in the current experimental paradigm, indoor scenes served as the frequent category and outdoor scenes as the infrequent category, without counterbalancing across participants. As such, condition-specific effects may reflect scene category encoding differences, rather than effects of stimulus category frequency. In addition, lower performance for frequent-category items created a restricted range of scores, potentially obscuring a usage–memory relationship in that condition. Although the aggregate predictor and exclusion rules were not preregistered, the primary memory associations were robust to the alternative exclusion choices examined here (Supplementary Table S3).

The current study focused on a restricted age cohort; future studies should investigate these relationships in developing adolescents and in clinical populations that may be more vulnerable to problematic or compulsive social media use. Longitudinal studies using logged-use behavioral telemetry, alongside randomized experimental designs, will be necessary to establish causal directionality, assess long-term cognitive trajectories and identify the psychological and neural mechanisms underlying the observed memory effects.

As short-form video becomes increasingly central to everyday information consumption and entertainment, understanding how rapidly shifting, algorithmically curated content relates to cognition is of growing importance. The present findings suggest that heavier short-form video use is associated with small but significant differences in episodic memory, particularly for infrequent-category items. These results provide a foundation for future work examining how short-form video use relates to learning, memory and cognitive engagement in daily life.

## Data availability

The data that support the findings of this study will be made publicly available upon journal publication.

## Code availability

Analysis code for this study will be made publicly available upon journal publication.

## Acknowledgements

We thank Krystal Li for participant data collection during initial experimental design, and Megan deBettencourt for sharing code and task materials.

## Author contributions

Conceptualization: K.K.C. and T.T. Methodology: K.K.C., D.S.M., A.D.W., and T.T. Formal analysis: K.K.C., D.S.M., and T.T. Investigation: K.K.C. Visualization: K.K.C. Writing–original draft: K.K.C. and T.T. Writing–review and editing: all authors. Supervision: A.D.W. and T.T.

## Funding

K.K.C. was supported by a Stanford Undergraduate Research Small Grant from the Office of the Vice Provost for Undergraduate Education. D.S.M. was supported in part by the Stanford University EDGE Doctoral Fellowship through the Office of the Vice Provost for Graduate Education (VPGE). T.T. was supported by a National Institute on Aging NRSA (F32AG071263), the Alzheimer’s Association (AARFD-21-852597), and NIH Grant P30AG066515.

The funders had no role in study design, data collection, analysis or interpretation, manuscript preparation, or the decision to submit the article for publication.

## Competing interests

The authors declare no competing interests.

## Declaration of generative AI and AI-assisted technologies in the manuscript preparation process

During manuscript preparation, the lead author used OpenAI Codex (GPT-6.1 Sol) and Devin (Cognition AI; Adaptive mode) to assist with language editing and consistency checks. Assistance with analysis and plotting scripts is described in the Methods. The lead author reviewed and edited the AI-assisted output. The authors take full responsibility for the manuscript.

## Supplementary Information

### Earlier local-sample study

Before the preregistered Prolific study, we conducted an earlier version of the SART-M paradigm in a locally recruited college-age sample (*N* = 338), composed primarily of Stanford University students and young adults from the surrounding Bay Area. This dataset informed task development and analysis planning. Participant characteristics and primary continuous associations are reported below. The continuous associations are visualized in Fig. S1.

**Table S1:** Participant characteristics for the local-sample study.

| **Characteristic** | Non-TikTok (*N* = 108) | TikTok (*N* = 230) |
|---|---|---|
| Age, yrs, mean ± SD | 20.1 ± 1.8 | 19.8 ± 1.7 |
| Sex, female, *N* (%) | 51 (47.2%) | 163 (70.9%) |
| Education, yrs, mean ± SD<sup>a</sup> | 14.6 ± 1.7 | 14.2 ± 1.7 |
| TikTok use, hr/wk, mean ± SD | 0.0 ± 0.0 | 16.8 ± 12.2 |
| **Race/Ethnicity** |  |  |
| Asian | 39 (36.1%) | 63 (27.4%) |
| White | 35 (32.4%) | 58 (25.2%) |
| More than one race | 14 (13.0%) | 33 (14.3%) |
| Other | 14 (13.0%) | 62 (27.0%) |
| Black/African American | 5 (4.6%) | 11 (4.8%) |
| Pacific Islander | 1 (0.9%) | – |
| Native American/Alaskan | – | 3 (1.3%) |

<sup>a</sup> Education summary statistics are based on participants with usable education data (*N* = 101 non-users, *N* = 198 TikTok users); all other rows use the full column samples.

**Table S2:** Continuous associations between TikTok use and cognitive performance in the local sample.

| Outcome | *b* per 1,000 min/week | *p* |
|---|---|---|
| Attention *d*′ | −0.07 | 0.14 |
| Overall memory *d*′ | −0.11 | < 0.001 |
| Frequent-category memory *d*′ | −0.05 | 0.10 |
| Infrequent-category memory *d*′ | −0.22 | < 0.001 |

Sex-adjusted ordinary least-squares models among TikTok users (*N* = 230). Coefficients are unstandardized slopes per 1,000 min/week; Fig. S1 uses the same fitted models.

**Figure S1:**
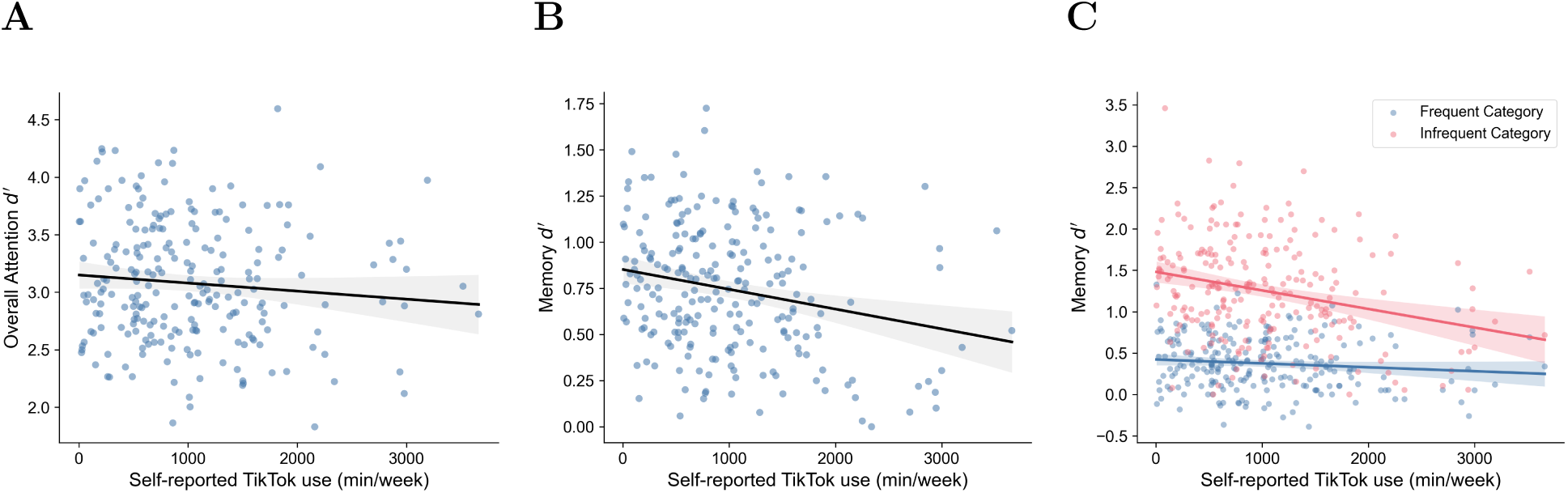
Continuous associations between self-reported TikTok use and cognitive performance in the earlier local pilot sample. Analyses included TikTok users with complete data (*N* = 230). **A**, Overall attention *d^′^*. **B**, Overall memory *d^′^*. **C**, Frequent-and infrequent-category memory *d^′^*. Points show participant observations; lines and 95% confidence bands show sex-adjusted model predictions at the sample mean of sex (Table S2). Use is expressed in min/week.

### Education coding

Education was used only descriptively. Three unambiguous formatting entries (e.g., “12+1”) were converted to numbers. Twelve responses were estimated: 10 were coded as 12 plus reported post-secondary years, provided the result did not exceed age minus 5, and 2 used qualification equivalents (some college, 14 years; post-graduate degree, 18 years). One response was left missing; other numerical self-reports were retained as reported. Original responses and coding decisions are provided in the codebook and education coding log.

### Sensitivity analyses for attention-accuracy exclusion threshold

We evaluated alternative attention-accuracy cutoffs of 2.0, 2.5 and 3.0 SDs below the mean, retaining the same exclusion sequence and sex adjustment (Table S3). Across all four thresholds, the primary memory associations remained significant and attention remained nonsignificant.

**Table S3:**
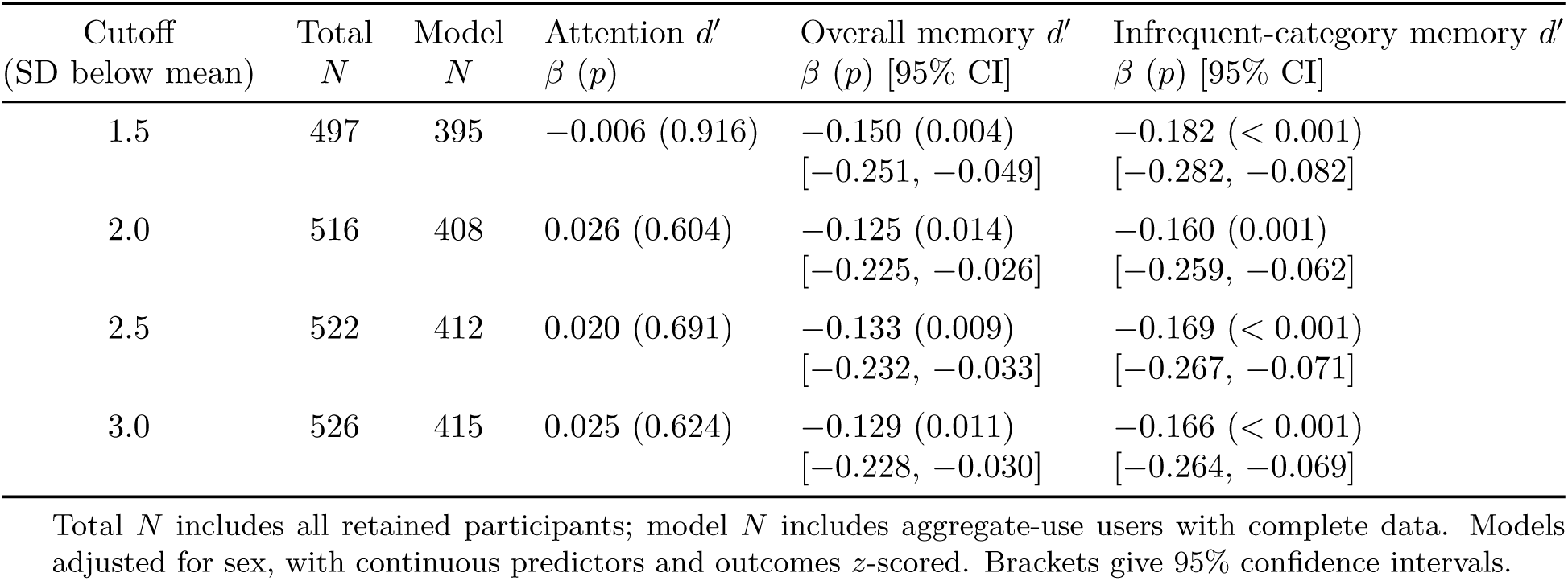
Sensitivity of primary findings to the attention-accuracy exclusion threshold.

| Cutoff | Total | Model | Attention *d*′ | Overall memory *d*′ | Infrequent-category memory *d*′ |
|---|---|---|---|---|---|
| (SD below mean) | *N* | *N* | β (*p*) | β (*p*) [95% CI] | β (*p*) [95% CI] |
| 1.5 | 497 | 395 | −0.006 (0.916) | −0.150 (0.004)<br>[−0.251, −0.049] | −0.182 (< 0.001)<br>[−0.282, −0.082] |
| 2.0 | 516 | 408 | 0.026 (0.604) | −0.125 (0.014)<br>[−0.225, −0.026] | −0.160 (0.001)<br>[−0.259, −0.062] |
| 2.5 | 522 | 412 | 0.020 (0.691) | −0.133 (0.009)<br>[−0.232, −0.033] | −0.169 (< 0.001)<br>[−0.267, −0.071] |
| 3.0 | 526 | 415 | 0.025 (0.624) | −0.129 (0.011)<br>[−0.228, −0.030] | −0.166 (< 0.001)<br>[−0.264, −0.069] |

Total *N* includes all retained participants; model *N* includes aggregate-use users with complete data. Models adjusted for sex, with continuous predictors and outcomes *z*-scored. Brackets give 95% confidence intervals.

Re-including participants excluded for negative overall memory *d^′^* (*N* = 16) or high aggregate use (*N* = 13) yielded *N* = 525 (422 aggregate-use users; one record with missing device-reported use was omitted). Associations remained negative for overall memory *d^′^*(*β* = *−*0.192, 95% CI [*−*0.289, *−*0.096], *p <* 0.001) and infrequent-category memory *d^′^* (*β* = *−*0.231, 95% CI [*−*0.326, *−*0.136], *p <* 0.001), but not frequent-category memory *d^′^* (*β* = *−*0.091, *p* = 0.067).

### Sensitivity to source-estimated weekly use

We refit the six primary models excluding participants whose weekly use included a daily-average estimate (*N* = 390 users). Overall memory *d^′^* (*β* = *−*0.162, *p* = 0.002) and infrequent-category memory *d^′^*(*β* = *−*0.199, *p <* 0.001) remained negatively associated with use; frequent-category memory *d^′^* (*β* = *−*0.044, *p* = 0.400) and attention outcomes remained nonsignificant.

### Sensitivity analyses using device-reported TikTok use alone

We refit the six primary sex-adjusted models using device-reported TikTok minutes alone, among TikTok users and all retained participants (exploratory analyses; Table S4).

**Table S4:**
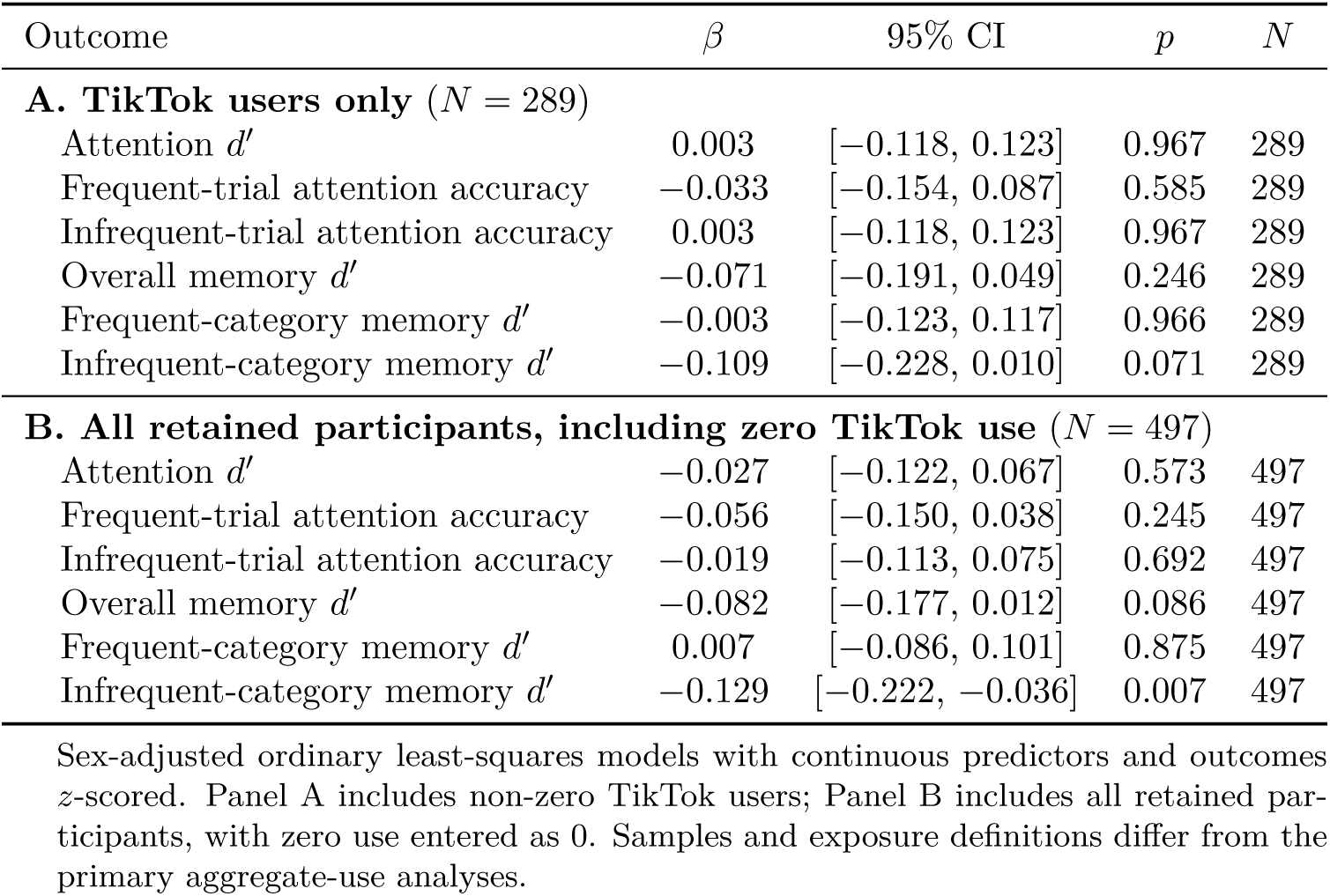
Sensitivity analyses using device-reported TikTok use alone.

### Exploratory platform-specific sensitivity analyses

We refit the six sex-adjusted models among aggregate-use users (*N* = 395), using TikTok minutes and total Instagram app minutes as separate predictors and including zero use (Table S5). TikTok use was negatively associated with infrequent-category memory; other platform-specific associations were nonsignificant.

**Table S5:** Exploratory platform-specific sensitivity analyses among aggregate-use users.

| Outcome | *β* | 95% CI | *p* | *N* |
|---|---|---|---|---|
| **A. TikTok minutes (device-reported)** (*N* = 395) |  |  |  |  |
| Attention *d*′ | −0.015 | [−0.120, 0.090] | 0.774 | 395 |
| Frequent-trial attention accuracy | −0.071 | [−0.176, 0.034] | 0.182 | 395 |
| Infrequent-trial attention accuracy | 0.011 | [−0.094, 0.116] | 0.844 | 395 |
| Overall memory *d*′ | −0.093 | [−0.197, 0.012] | 0.082 | 395 |
| Frequent-category memory *d*′ | 0.001 | [−0.103, 0.105] | 0.990 | 395 |
| Infrequent-category memory *d*′ | −0.131 | [−0.234, −0.027] | 0.014 | 395 |
| **B. Total Instagram app use (device-reported)** (*N* = 395) |  |  |  |  |
| Attention *d*′ | 0.014 | [−0.086, 0.115] | 0.778 | 395 |
| Frequent-trial attention accuracy | 0.036 | [−0.064, 0.137] | 0.477 | 395 |
| Infrequent-trial attention accuracy | −0.006 | [−0.106, 0.094] | 0.907 | 395 |
| Overall memory *d*′ | −0.092 | [−0.192, 0.008] | 0.071 | 395 |
| Frequent-category memory *d*′ | −0.066 | [−0.166, 0.033] | 0.190 | 395 |
| Infrequent-category memory *d*′ | −0.084 | [−0.184, 0.015] | 0.096 | 395 |

Standardized, sex-adjusted ordinary least-squares models among the 395 aggregate-use users, including zero use of the platform being analyzed. Instagram minutes reflect total app use, including non-Reels activity.

### Additional analyses in the Prolific sample

**Figure S2:**
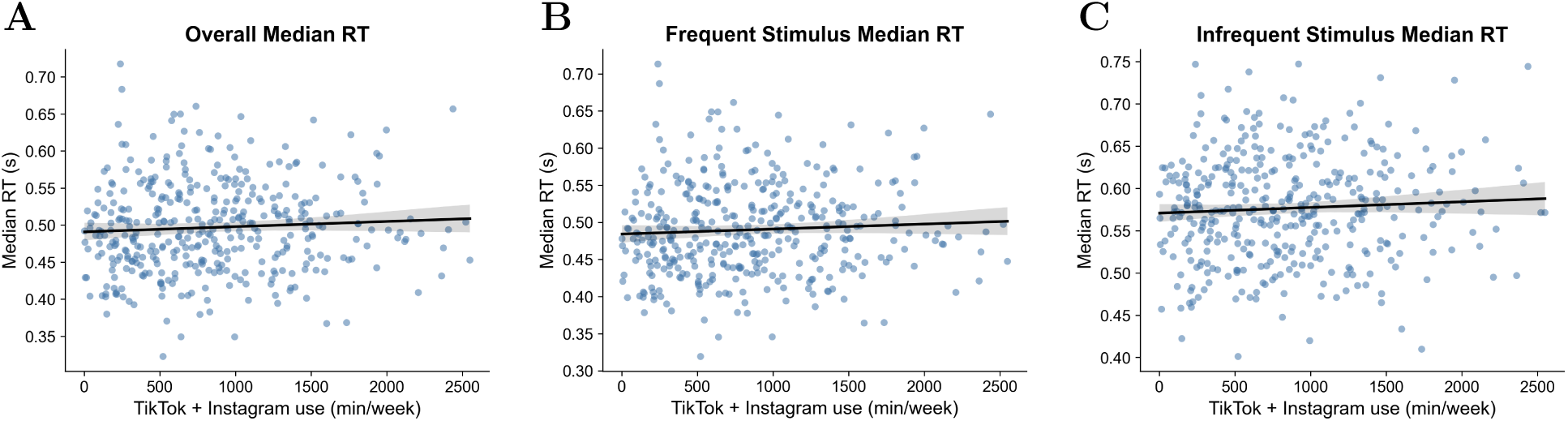
Median reaction time vs. aggregate short-form video use. Participant-level median RT in s for overall (**A**; *β* = 0.06, *p* = 0.25), frequent-trial (**B**; *β* = 0.06, *p* = 0.26), and infrequent-trial (**C**; *β* = 0.04, *p* = 0.40) responses. Lines show unadjusted ordinary least-squares fits with 95% bootstrap confidence bands; reported statistics are sex-adjusted (*N* = 395).

**Figure S3:**
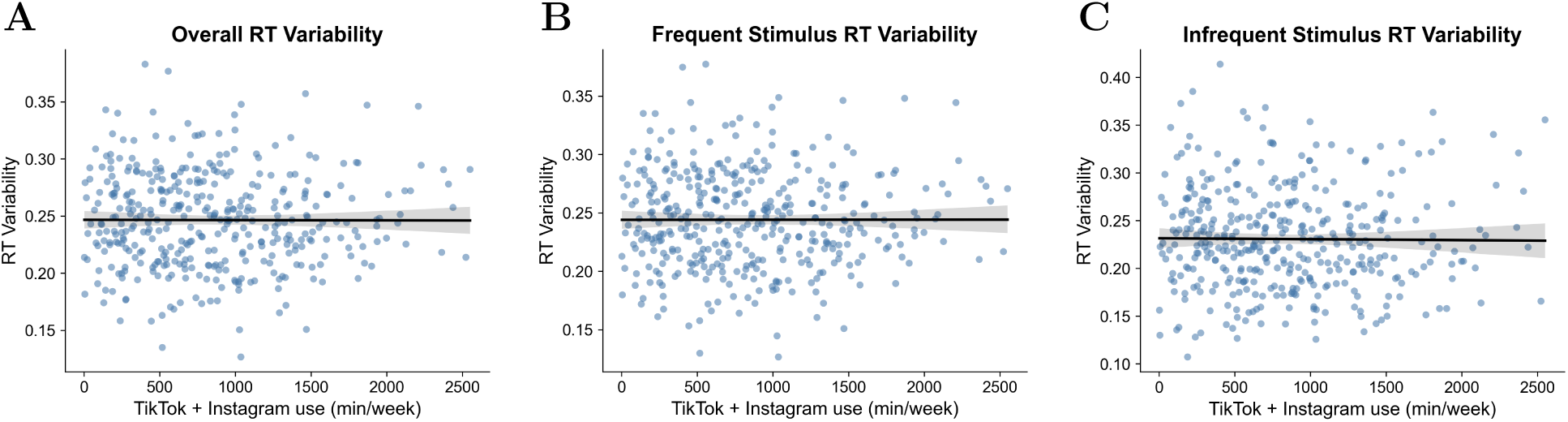
RT variability vs. aggregate short-form video use. Participant-level RT coefficients of variation (population SD/mean) for overall (**A**; *β* = *−*0.03, *p* = 0.55), frequent-trial (**B**; *β* = *−*0.03, *p* = 0.58), and infrequent-trial (**C**; *β* = *−*0.02, *p* = 0.69) responses. Lines show unadjusted ordinary least-squares fits with 95% bootstrap confidence bands; reported statistics are sex-adjusted (*N* = 395).

**Figure S4:**
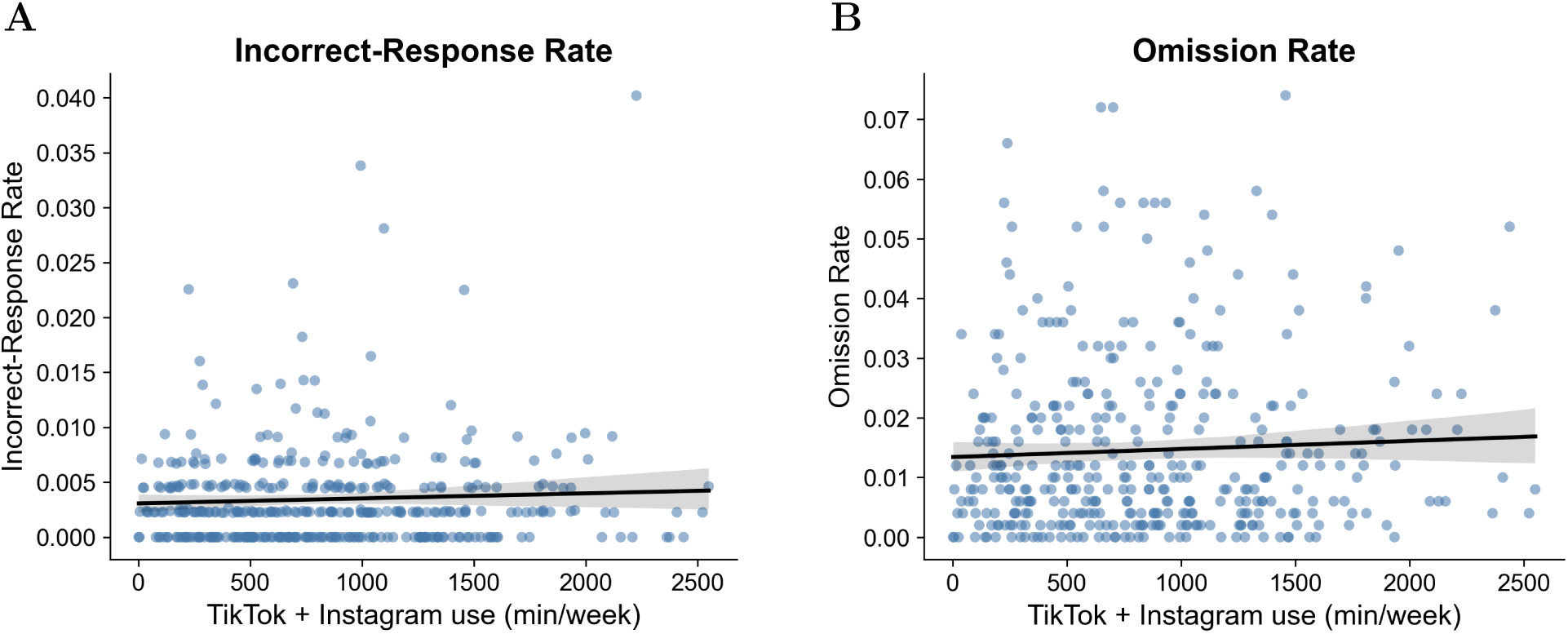
SART incorrect-response and omission rates vs. aggregate short-form video use. **A**, Frequent-trial incorrect-response rate (*β* = 0.05, *p* = 0.38). **B**, Omission rate (*β* = 0.03, *p* = 0.54). Points show participant-level rates. Lines show unadjusted ordinary least-squares fits with 95% bootstrap confidence bands; reported statistics are sex-adjusted (*N* = 395).

**Figure S5:**
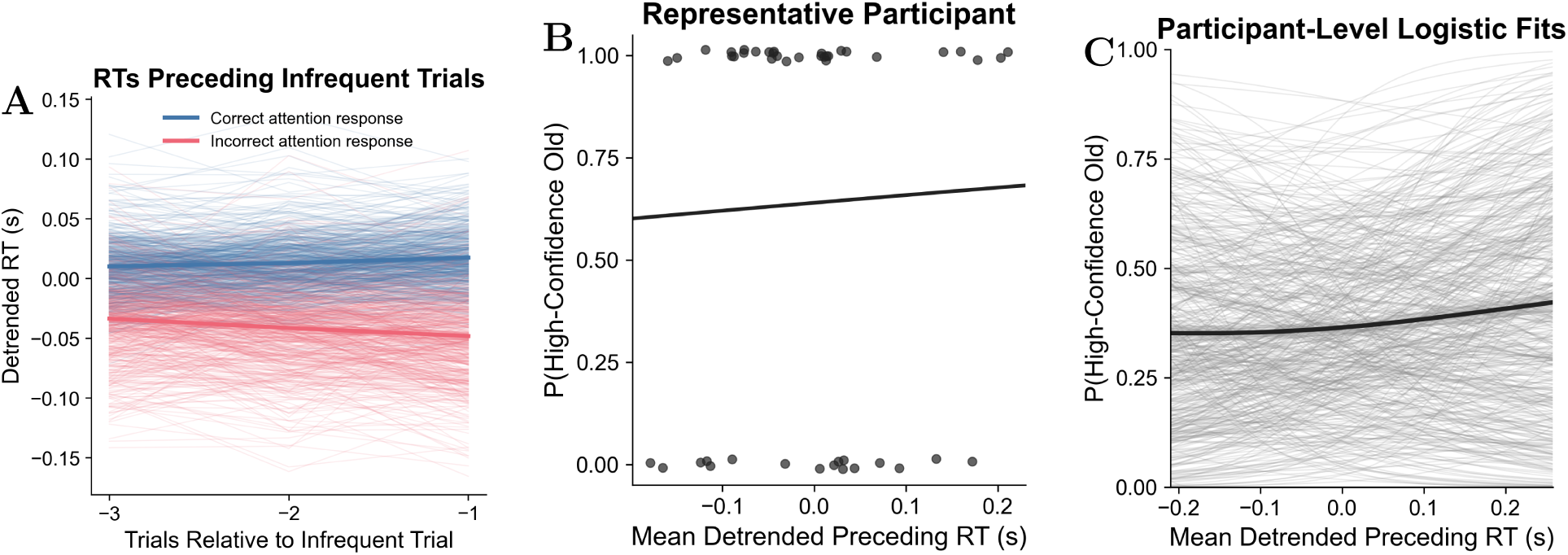
**Exploratory adaptation of the sustained-attention/subsequent-memory analysis of deBettencourt et al.**^13^ We tested whether mean detrended RT across the three trials preceding each infrequent SART trial predicted subsequent high-confidence memory (response 4; primary analyses used responses 3–4). **A**, Preceding RTs grouped by correct/incorrect infrequent attention responses. **B**, Example participant-level logistic fit with slope closest to the sample median. **C**, Participant-level logistic fits. Across 496 estimable participants, the mean slope was positive (0.51 per s, 95% bootstrap CI [*−*0.12, 1.03]) but nonsignificant (two-sided participant-resampling *p* = 0.103; directional one-sided *p* = 0.051).

## Notes

### Competing Interest Statement

The authors have declared no competing interest.

### Summary of Updates

This revision updates data processing and questionnaire scoring, documents repeat-attempt handling and exclusions, and revises the final analytic sample from 502 to 497 participants. Sensitivity analyses and clarifications of the reported methods and changes from the preregistration have been added, with corresponding updates to results, figures and analysis code. The main association between greater device-reported use and poorer episodic memory remains, while sustained-attention associations remain nonsignificant.

